# Anchorene is an endogenous diapocarotenoid required for anchor root formation in Arabidopsis

**DOI:** 10.1101/496737

**Authors:** Kun-Peng Jia, Alexandra J. Dickinson, Jianing Mi, Guoxin Cui, Najeh M. Kharbatia, Xiujie Guo, Erli Sugiono, Manuel Aranda, Magnus Rueping, Philip N. Benfey, Salim Al-Babili

## Abstract

Arabidopsis root development is predicted to be regulated by yet unidentified carotenoid-derived metabolite(s). In this work, we screened known and putative carotenoid cleavage products and identified anchorene, a predicted carotenoid-derived dialdehyde (diapocarotenoid) that triggers anchor root development. Anchor roots are the least characterized type of root in Arabidopsis. They form at the root-shoot junction, particularly upon damage to the root apical meristem. Using Arabidopsis reporter lines, mutants and chemical inhibitors, we show that anchor roots originate from pericycle cells and that the development of this root type is auxin-dependent and requires carotenoid biosynthesis. Transcriptome analysis and treatment of auxin-reporter lines indicate that anchorene triggers anchor root development by modulating auxin homeostasis. Exogenous application of anchorene restored anchor root development in carotenoid-deficient plants, indicating that this compound is the carotenoid-derived signal required for anchor root development. Chemical modifications of anchorene led to a loss of anchor root promoting activity, suggesting that this compound is highly specific. Furthermore, we demonstrate by LC-MS analysis that anchorene is a natural, endogenous Arabidopsis metabolite. Taken together, our work reveals a new member of the family of carotenoid-derived regulatory metabolites and hormones.

**Significance:** Unknown carotenoid-derived compounds are predicted to regulate different aspects of plant development. Here, we characterize the development of anchor roots, the least characterized root type in Arabidopsis, and show that this process depends on auxin and requires a carotenoid-derived metabolite. We identified a presumed carotenoid-derivative, anchorene, as the likely, specific signal involved in anchor root formation. We further show that anchorene is a natural metabolite that occurs in Arabidopsis. Based on the analysis of auxin reporter lines and transcriptome data, we provide evidence that anchorene triggers the growth of anchor roots by modulating auxin homeostasis. Taken together, our work identifies a novel carotenoid-derived growth regulator with a specific developmental function.

## Introduction

Carotenoids are common isoprenoid pigments synthesized by all photosynthetic organisms and many heterotrophic bacteria and fungi (1–4). They are essential constituents of the photosynthetic apparatus (5), as well as sources for biologically important compounds such as the retinoids (6) and the phytohormones abscisic acid (ABA) (7) and strigolactone (SL) (8). All of these derivatives arise by virtue of the extended, conjugated double bond system that makes carotenoids prone to oxidative cleavage (9). This reaction yields carbonyl products called apocarotenoids (2, 3) and can be catalyzed by enzymes from the carotenoid cleavage dioxygenase (CCD) family, which break defined C-C double bonds by inserting molecular oxygen (9). Arabidopsis CCDs are divided into nine-*cis*-epoxycarotenoid cleavage dioxygenases (NCED2, 3, 5, 6, and 9) that form the ABA precursor xanthoxin from 9-*cis*-epoxycarotenoids and CCDs with different substrate and regio-specificity (9, 10). The latter group includes CCD1, which forms a plentitude of C_13_, C_10_ and C_8_ volatiles from different apocarotenoids and C_40_-carotenoids (11), CCD4, which cleaves all-*trans*-β-carotene into β-ionone (C_13_) and β-apo-10’-carotenal (C_27_) (12), the SL biosynthesis enzyme CCD7 (MAX3), which breaks 9-*cis*-β-carotene into β-ionone (C_13_) and 9-*cis*-β-apo-10’-carotenal that is converted by CCD8 (MAX4) into the SL biosynthesis intermediate carlactone (8). CCD8 can also cleave all-*trans*-β-apo-10’-carotenal into the ketone β-apo-13-carotenone (d’orenone), but with low activity (13). Non-enzymatic cleavage, which occurs at each double bond in the carotenoid backbone, is another important route for apocarotenoid formation (14). This process is triggered by reactive oxygen species (ROS) and can also yield signaling molecules, such as the plant stress signal β-cyclocitral, which is formed by singlet oxygen (^1^O_2_) and mediates gene responses to this ROS (15). In addition to monocarbonyls, carotenoid cleavage can yield dialdehyde products (diapocarotenoids), as shown for several plant and cyanobacterial CCDs that cleave multiple double bonds within carotenoids or target apocarotenoids (16, 17). The question of whether diapocarotenoids are also regulatory metabolites has not yet been answered.

Plant root systems provide anchorage and are the primary site for water and nutrient uptake (18, 19). Arabidopsis is an ideal model plant to study root development because of its genetic tractability and its fast-growing and relatively simple root system. Arabidopsis has three highly characterized types of roots: i) a primary root initiated in embryogenesis, ii) lateral roots (LRs) that form from the primary root and other LRs, and iii) adventitious roots, which emerge from non-root tissues, such as stem, leaves and hypocotyl (18, 19). Anchor roots (ANRs) (20) constitute a fourth type of roots. They emerge from the collet (21), a region located just below the root-hypocotyl junction, and have remained largely uncharacterized.

The development of LRs occurs in a series of well-documented stages. They are positioned through a process that involves gene expression oscillation (22) mediated by an unidentified carotenoid-derived signal (23). In Arabidopsis, LRs initiate from xylem pole pericycle cells, which form the cell layer between the vascular bundle and the endodermis. These cells divide to produce LR primordia, which continue to grow until the new roots emerge by pushing through the outer layers of the originating root (24). The plant hormone auxin plays a central role in root development (25, 26). For instance, the auxin signaling component *MP/ARF5* regulates embryonic root development (27) and the auxin-related transcription factors *ARF7* and *ARF19* are pivotal for LR initiation (28).

In addition to auxin, the carotenoid-derived phytohormones, SL and ABA, have been shown to regulate different aspects of root development (29, 30). To identify novel carotenoid-derived compounds involved in Arabidopsis root development, we tested the activity of apocarotenoids and diapocarotenoids, which had either been previously identified as CCD products of or were structurally predicted to result from carotenoids. We discovered that ANR formation requires carotenoid biosynthesis and is triggered by a novel diapocarotenoid that we called “anchorene”. To characterize the role of anchorene, we present the first comprehensive analysis of ANR development.

## Results

### Anchorene is a specific regulator of ANR development

To identify new carotenoid-derived signals involved in root development, we checked the activity of 6 previously identified or predicted diapocarotenoids with chain lengths ranging C_9_ to C_15_ (Diapo1-6; for structures, see Fig. S1A and S1B). These compounds were selected because they are available and stable at ambient conditions. Diapo1 (C_9_) is the expected product formed upon CCD8 cleavage of all-*trans*-β-apo-10’-carotenal (13). Diapo2 (C_10_) results from cleaving the C7, C8 and C15, C15’ double bonds in different carotenoids and is produced by cyanobacterial retinal-forming enzymes from C_30_-apocarotenoids (17). Diapo3 (C_10_) is a structural isomer of Diapo2 and a predicted cleavage product that can be formed by cutting the C11, C12 and C11’, C12’ bonds in almost all plant carotenoids (Fig. 1A and Fig. S2). Diapo4 (C_12_) results from cleaving the C7, C8 and C13’, C14’ bonds in many carotenoids, and is formed by CCD1 from apo-10’-lycopenal (20). Diapo5 (C_15_) results from cleaving the C7, C8 and C11’, C12’ bonds in many apocarotenoids and is formed by CCD1 from apo-10’-lycopenal (20). Diapo6 (C_14_) is a common CCD1 product formed by cleaving the C19, C10 and C9’, C10’ bonds in many carotenoids (16). We treated seedlings with either 5 or 25 μM of each compound, and determined the length of primary roots 7 days post-stratification (dps). At lower concentrations, we did not observe significant effects with any of the compounds. At the higher concentration, application of Diapo2 and Diapo5 led to a severe decrease (approximately 80%) in primary root growth (Fig. S3A and S3B). Diapo1, Diapo3 and Diapo4 showed weak inhibition of primary root length, however, the most striking effect of Diapo3 was the promotion ANR formation (Fig. S3A and S3B). Based on this activity, we named Diapo3 “anchorene.”

**Fig. 1.**
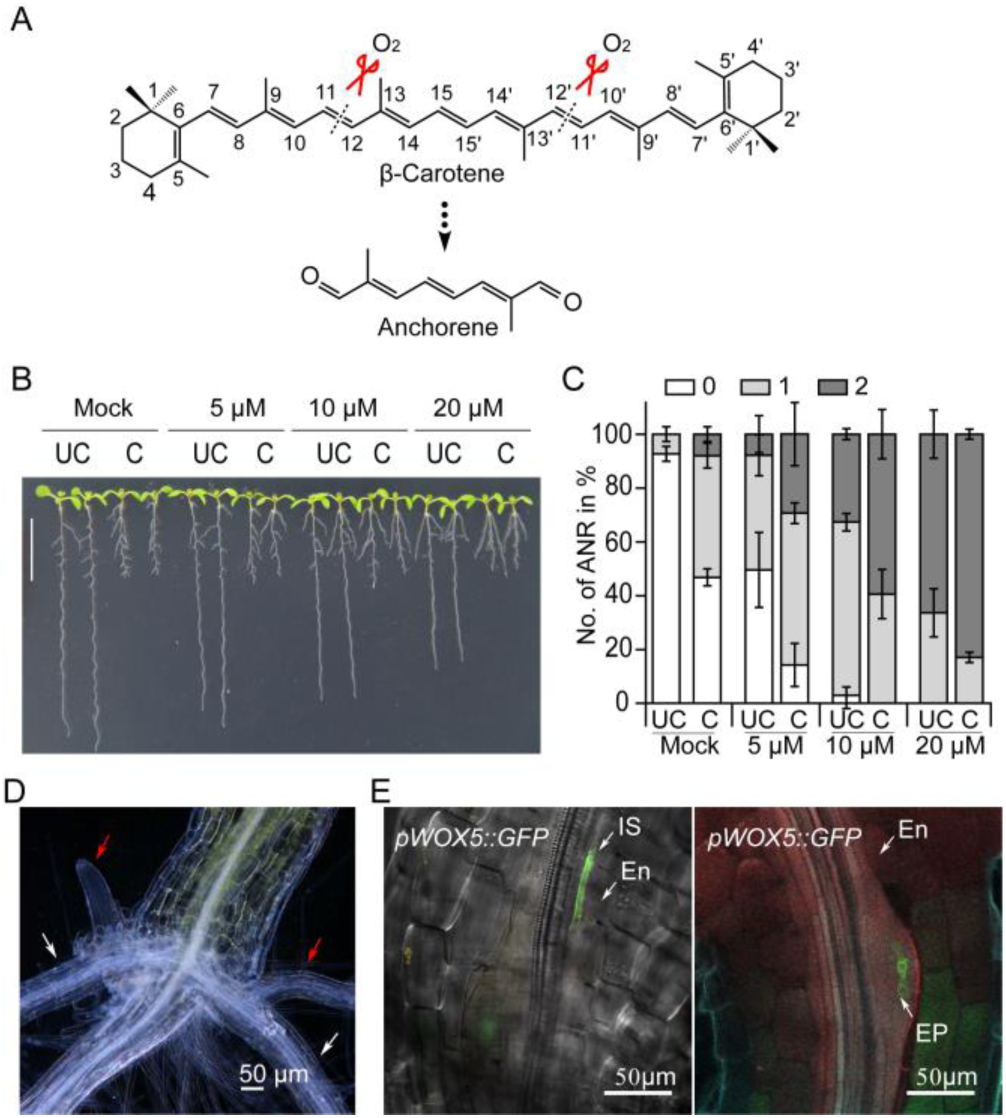
Anchorene promotes anchor root formation. (A) Structure of anchorene. (B) Representative seedlings show that anchorene specifically promotes ANR formation in a concentration dependent manner. Scale bar is 1 cm. (C) Quantification of the effect of anchorene on ANR formation. Data are presented as percentage (mean ± SD) of seedlings with 0, 1 or 2 ANRs from three independent replicates. In B and C, “UC” indicates uncut RAMs, “C” indicates cut RAMs. (D) Two opposing ANRs are formed at the collet. White arrows indicate primary ANRs, and red arrows indicate secondary ANRs (or LRs). (F) ANRs are initiated from the pericycle cells of root; *pWOX5::GFP* was used to indicate the ANR initiation by confocal microscopy imaging. “IS” indicates the initiation site; “En” indicates the endodermis; “EP” indicates the emerging primordia; AR: anchorene.

Excision of the root apical meristem (RAM) triggers ANR formation (20) (Fig. S3C and S3D). To exclude the possibility that anchorene’s effect on ANR formation is a result of the weak inhibition of primary root growth (Fig. S3E), we applied different concentrations of anchorene to roots with or without RAM excision. About 9% of control Col-0 seedlings developed ANRs under normal conditions, compared to 50% upon RAM excision (Fig. 1B and 1C). The effect of 5 μM anchorene was comparable to that of RAM excision, triggering the formation of ANRs in 55% of the seedlings. Higher anchorene concentrations (10 and 20 µM) enhanced this ratio to 97% and 100%, respectively (Fig. 1B and 1C). There was also an increase in the number of seedlings that developed two ANRs from 0% in control to approximately 80% upon application of 20 μM anchorene (Fig. 1B and 1C). Using a wide range of concentrations, we established a dose-response curve in the presence and absence of RAM excision. The effect of anchorene was dose dependent in both cases (Fig. S3F). Taken together, our results show that anchorene triggers ANR formation regardless of RAM excision and does not affect LR formation (Fig. S3G), which demonstrates that its promotion of ANRs is not caused by inhibiting primary root growth.

To test the specificity of anchorene, we evaluated the effect of structurally similar compounds on ANR formation. The application of Diapo2, a structural isomer of anchorene (Fig. S1B), gave only a modest increase in ANR formation, even at the relatively high concentration of 25 μM (Fig. S3H and S3I). However, this concentration caused a pronounced inhibition of primary root growth, which may be the reason for the stimulatory effect on ANR formation. Modification of anchorene’s structure by reducing the aldehyde groups to alcohols or converting the aldehydes into acids or acid-ethyl esters resulted in a loss of activity (Fig. S1C, Fig. S3H and S3I). These data suggest that anchorene promotion of ANR formation requires specific structural features found only in anchorene, among the compounds tested.

To understand how anchorene exerts its activity, we characterized its effect on ANR development using stereo and confocal microscopy. ANRs arise in the collet region characterized by the presence of dense root hairs (21) (Fig. 1D). Arabidopsis seedlings form one or two ANRs opposite each other, mirroring the positions of the cotyledons (Fig. S3J). In contrast, LRs emerge in much higher numbers at alternating positions (31). ANRs themselves can branch, forming secondary ANRs or LRs (Fig. 1D). The cellular pattern of ANR primordia in the collet indicates that they originate, similar to LRs, from the xylem pole pericycle. Analysis of GFP signals in a *pWOX5::GFP* marker line (32), which is induced upon formation of LR primordia, confirmed the pericycle origin of ANR primordia (Fig. 1E).

To track ANR development, we used a pDR5::LUC line, which marks LR prebranch sites (22, 23). We observed a clear LUC signal in the collet as early as 3 dps (Fig. S4A), suggesting that this line also visualizes ANR primordia. Consistent with anchorene effect on ANR formation, the application of this compound intensified the LUC signal in the collet (Fig. S4B). Similarly, anchorene application to *pDR5rev::GFP* seedlings led to a strong GFP signal at the same site (Fig. S4C). In anchorene-treated *pDR5rev::GFP* seedlings, ANR primordia initiation and ANR emergence were observed at 3 dps and 6 dps, respectively (Fig. S4C).

### A carotenoid-derived metabolite is required for normal ANR formation

To understand the role of carotenoids in ANR development, we monitored ANR formation under carotenoid-deficient conditions. Arabidopsis seedlings grown on media frequently do not form emerged ANRs, which impeded the characterization of factors affecting ANR formation. Therefore, we monitored ANR formation after RAM excision (ANR-RE), in order to stimulate ANR development (Fig. S3C and S3D) (20). We used these measurements as an indicator of ANR formation capacity. We first measured ANR formation upon treatment with norflurazon (NF) and 2-(4-chlorophenylthio)-triethylamine hydrochloride (CPTA) (4), which block phytoene desaturation and lycopene cyclization, respectively (Fig. 2A). We also investigated the effect of D15, a CCD inhibitor (23). These three compounds have been shown to reduce LR initiation (23) (Fig. 2B and Fig. S5), suggesting that a CCD product is required for LR development. Interestingly, both NF and CPTA strongly reduced ANR-RE, while D15 promoted this process (Fig. 2B and Fig. S5). These results suggest that carotenoids are necessary for proper ANR and LR development, but that each root type is regulated by a specific apocarotenoid. To test this hypothesis, we asked if anchorene can restore LR capacity in D15-treated seedlings. Anchorene had no significant effect on LR formation after D15 treatment. Furthermore, it inhibits LR capacity in untreated seedlings (Fig. S6), corroborating evidence that anchorene is not the carotenoid-derived signal required for LR capacity. The positive effect of D15 on ANR-RE may be caused by increased carotenoid levels or by its inhibitory effect on primary root growth (23). Confirming the carotenoid-dependency of ANR formation, the carotenoid deficient mutants, *ispH1* (33) and *psy* (34), respectively disrupted in plastid isoprenoid and phytoene biosynthesis (Fig. 2A), exhibited greatly reduced ANR-RE, compared to wild-type (Fig. 2C and 2D).

**Fig. 2.**
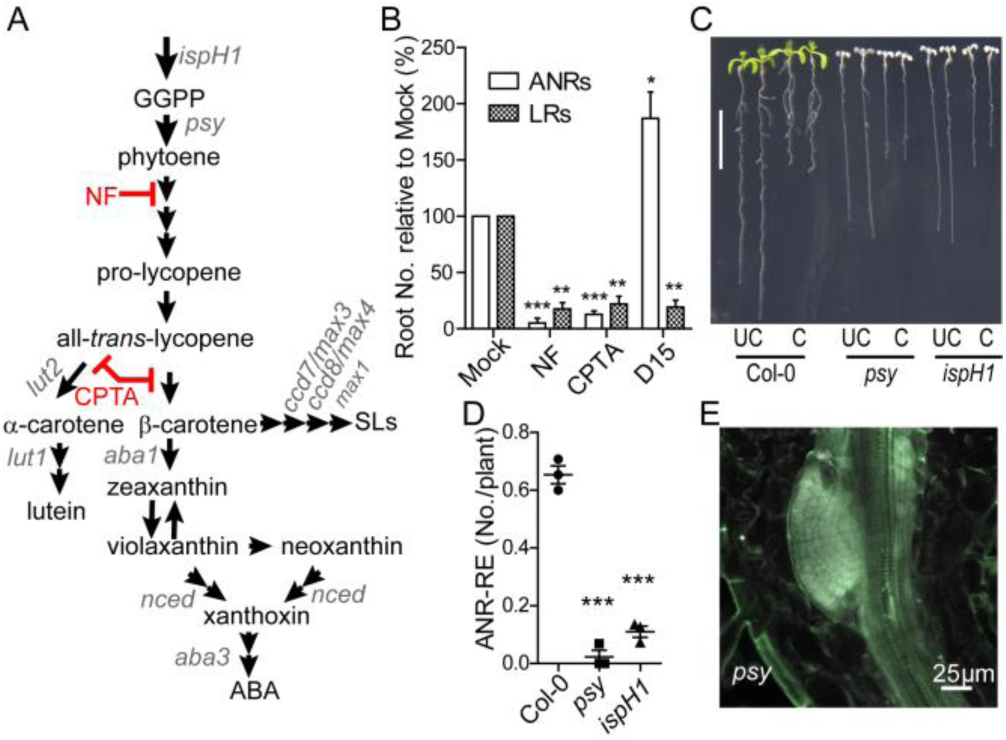
Carotenoid-deficient seedlings have reduced ANR-RE (ANR formation after RAM excision). (A) Schematic of plant carotenoid biosynthesis. Mutants are shown in gray italics. Reactions inhibited by NF and CPTA are depicted in red. (B) Quantification of ANRs and LRs formation after RAM excision in Mock, NF-, CPTA-and D15-treated Col-0 seedlings. Data are presented as mean ± SD from three independent replicates; two-tailed paired Student’s *t*-tests were performed in comparison to corresponding Mock treatments (\**P* <0.05; \*\**P* <0.01; \*\*\**P* <0.001). (C) Representative seedlings show the ANR formation in Col-0, *psy* and *ispH1*. “UC” indicates uncut RAMs; “C” indicates cut RAMs. (D) ANR-RE quantification in Col-0, *psy* and *ispH1* seedlings. Data are presented as mean ± SD from three independent replicates; two-tailed paired Student’s *t*-test (\*\*\**P* <0.001). (E) A representative *psy* seedling has initiated ANR primordia. 5 dps seedlings were used for confocal imaging.

CPTA inhibits the synthesis of both α- and β-carotene, which mark the two branches of plant carotenoid biosynthesis (35) (Fig. 2A). Measurement of ANR-RE in *lut1* (36) and *lut2* (37), two mutants respectively deficient in α-carotene and lutein formation, did not reveal a difference compared to wild-type (Fig. S7A). This result indicates that the α-branch is not the primary source for the ANR signal. Next, we asked whether ABA or SL is the apocarotenoid signal required for ANR formation. We observed an increase in ANR-RE in the SL deficient *ccd8/max4* (38) and *max1* (39) mutants and an inhibitory effect of the SL analog GR24 on ANR-RE (Fig. S7B, S7C), indicating a negative role of SL in ANR formation, although we didn’t see a significant ANR-RE change in *ccd7/max3* seedlings. We did not detect a change in ANR-RE in the ABA deficient mutants, *aba1* (40) and *aba3* (41), or upon ABA application (Fig. S7D and S7E), excluding a role of ABA in ANR development. We also determined ANR-RE in *nced* and *ccd* mutants, i.e. *nced2, nced3, nced5, nced6, nced9, ccd1*, and *ccd4*, but did not observe a significant change in ANR-RE (Fig. S7F), indicating that the corresponding enzymes are unnecessary or work redundantly in this regard.

To determine at which stage the apocarotenoid signal regulates ANR development, we examined the effect of NF on the *pDR5::LUC* marker line. In NF treated plants, we detected a clear collet LUC signal (Fig. S4B), suggesting that carotenoids are not required for ANR initiation. This assumption is corroborated by the presence of ANR primordia in carotenoid-deficient *psy* mutant seedlings (Fig. 2E).

### Anchorene is an endogenous signal that rescues ANR formation under carotenoid deficient conditions

To determine if anchorene is the carotenoid-derived metabolite required for ANR formation, we asked if it could rescue the ANR-RE reduction caused by inhibiting carotenoid biosynthesis. Indeed, seedlings co-treated with anchorene and either NF or CPTA did not have decreased ANR-RE formation compared to untreated seedlings (Fig. 3A and Fig. S8A). Similarly, anchorene treatment restored wild-type ANR-RE in the *psy* mutant (Fig. 3B and Fig. S8B). These results indicate that anchorene is sufficient to promote ANR formation under carotenoid-deficient conditions and also exclude the possibility that reduction in ANR-RE observed in carotenoid deficient seedlings is an indirect consequence of albinism. We also examined the effect of NF alone and in combination with anchorene on *pDR5rev::GFP* seedlings. We did not see a significant reduction in GFP signal after NF application, however, the combined NF/anchorene treatment clearly enhanced the GFP signal (Fig. S9), suggesting that anchorene stimulated growth of ANR primordia post-initiation.

**Fig. 3.**
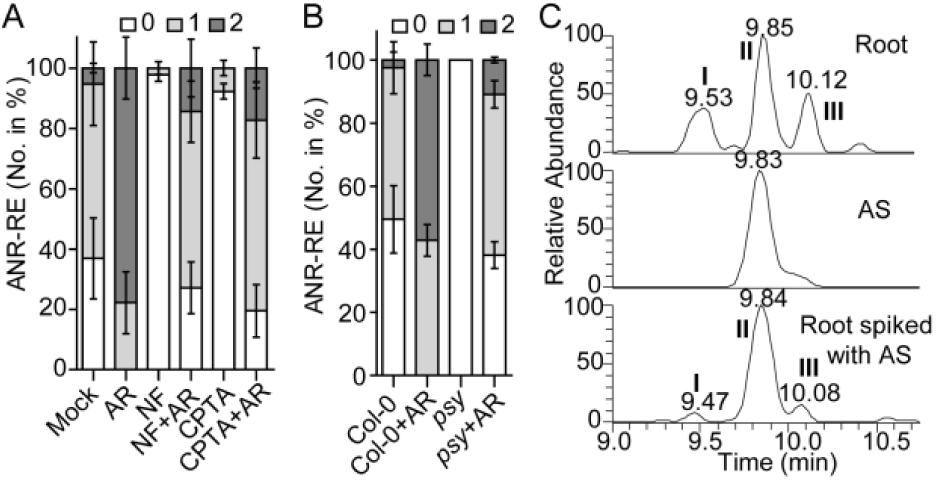
Anchorene (AR) is an endogenous metabolite restoring ANR-RE in carotenoid deficient seedlings. (A) Quantification of ANR-RE in NF-and CPTA-treated seedlings exposed to anchorene. (B) Quantification of ANR-RE in *psy* mutant seedlings treated with anchorene. In A and B, data are presented as the percentage (mean ± SD) of seedlings with 0, 1 or 2 ANRs from three independent replicates. 1 µM NF, 100 µM CPTA and 20 µM anchorene were used. (C) LC-MS identification of endogenous anchorene in Arabidopsis roots extract. Extracted ion chromatograms (EICs) of anchorene from Arabidopsis root extract (upper), anchorene standard (middle) and Arabidopsis root extract spiked with anchorene standard (bottom). Peak II indicates endogenous anchorene (upper) or endogenous anchorene spiked with anchorene standard (bottom). Peaks I and III represent anchorene isomers, according to their accurate mass and product ion spectra (Fig. S10B). (D) Endogenous anchorene (peak II) from Arabidopsis extract and anchorene standard displayed identical pattern of product ion spectra. AS: anchorene standard.

To determine if anchorene is a natural plant metabolite, we developed an extraction, derivatization and Liquid Chromatography-Mass Spectrometry (LC-MS) protocol for carotenoid-derived diapocarotenoids. We identified endogenous anchorene from Arabidopsis root and shoot tissues based on its precise match with the mass, chromatographic retention time and product ion spectra of the authentic anchorene standard (Fig. 3C, Fig. S10A and S10B). Anchorene content in shoot tissues was about 4-fold higher than in root tissues (0.08±0.003 versus 0.02±0.00 1 pmol/mg dry weight) (Fig. S10C). We also observed two potential anchorene isomers identified by their exact mass and product ion spectra (Fig. 3C and Fig. S10B). Interestingly, the relative amounts of the three isomers differ between root and shoot samples, with the anchorene peak being the most pronounced peak in the root (Fig. S10A). This pattern is consistent with anchorene’s role in root development. Moreover, we quantified the anchorene content after NF treatment to confirm its carotenoid origin. Consistently, the anchorene content of the whole seedlings decreases after 24 hours NF treatment, and greatly decreases in NF continuous treated seedlings (Fig. S10D and S10E).

Anchorene could be produced from the cleavage of C11-C12 and C11’-C12’ double bonds of most carotenoids. To further confirm its carotenoid origin and to test its possible precursor, we fed the Arabidopsis seedlings with OH-APO10’ and OH-APO12’ (Fig. S12A) and then measured the anchorene content. Interestingly, OH-APO12’ fed Arabidopsis seedlings contained much more anchorene compared to control and OH-APO10’ fed seedlings (Fig. S12B). Furthermore, OH-APO12’ but not OH-APO10’ treated Arabidopsis seedlings formed more ANRs compared to mock (Fig. S12C). This suggests that OH-APO12’ is likely acting as the direct precursor to produce anchorene (Fig. S12D).

### Anchorene promotes ANR formation by modulating auxin distribution

Anchorene’s promotion of signal in the DR5 marker line suggested the involvement of auxin in ANR formation. Therefore, we investigated the role of ARF7 and ARF19, two auxin responsive transcription factors required for LR initiation (26, 28), in this process. The *arf7arf19* double mutant did not form any ANRs after anchorene application or RAM excision (Fig. 4A and Fig. S11A). Moreover, confocal microscopy revealed that *arf7arf19* lacks ANR primordia (Fig. 4B), suggesting that *ARF7* and *ARF19* are crucial for ANR initiation. Next, we examined the effect of the auxin analog 1-Naphthaleneacetic acid (NAA) and the auxin transport inhibitor 1-N-naphthylphthalamic acid (NPA). NAA greatly increased ANR-RE, while NPA treatment completely blocked ANR-RE (Fig. 4C and Fig. S11B). These results demonstrate that auxin signaling is required for ANR initiation and that auxin transport is essential for ANR formation.

**Fig. 4.**
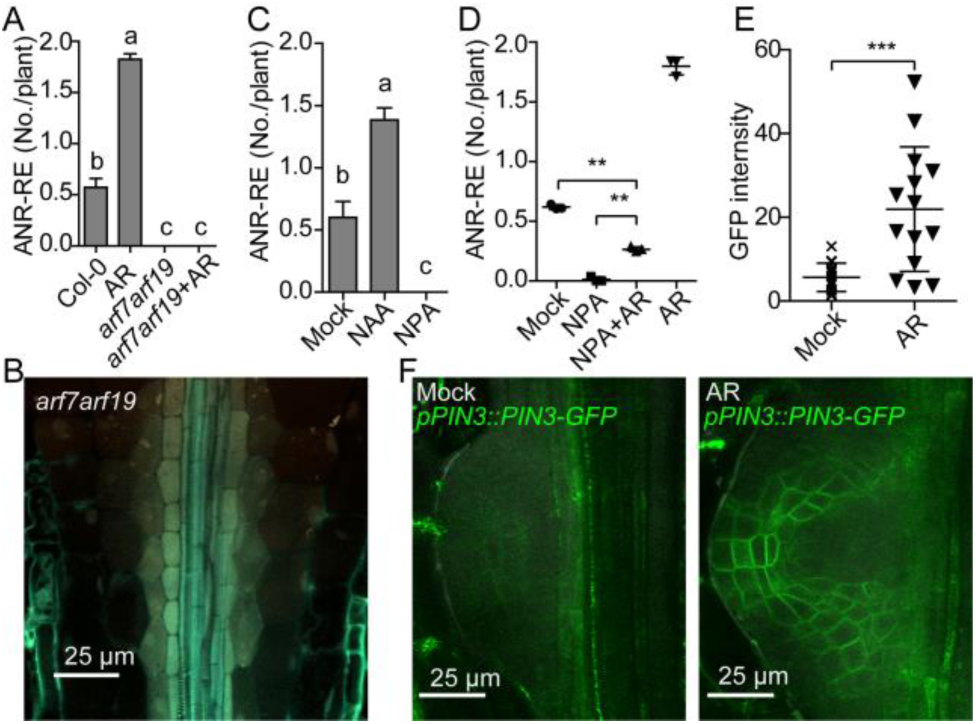
Role of auxin signaling and distribution in ANR development. (A) Effects of anchorene on ANR-RE in Col-0 and *arf7arf19* seedlings. (B) *arf7arf19* has no ANR primordial, as shown by confocal microscope imaging. (C) Effect of auxin analog NAA and auxin transport inhibitor NPA on ANR-RE. In A and C, data are presented as mean ± SEM from one representative experiment; different letters denote significant differences (one-way ANOVA with Tukey multiple-comparison test, *P* < 0.05); in A, n=49, 51, 39, 43, 44, 50 individually, in C, n=25, 26, 28, 25 individually. (D) Anchorene partially rescued ANR-RE in NPA treated seedlings. Data are presented as mean ± SD from three independent replicates; two-tailed paired Student’s *t*-test (\*\**P* <0.01). (E) Quantification of anchorene effect on PIN3-GFP fluorescent intensity in ANR primordia. Data are presented as mean ± SD (Two tailed Student’s *t*-test, \*\*\**P* <0.001); n=15, 13 respectively, from two independent experiments. (F) PIN3-GFP expression in ANR primordia of mock and anchorene treated seedlings shown by confocal microscope imaging. 0.1 µM NAA, 1 µM NPA and 20 µM anchorene were used. AR: anchorene.

To determine if anchorene acts by modulating auxin transport, we evaluated its effect in the presence of NPA. Anchorene partially rescued the negative impact of NPA on ANR-RE (Fig. 4D and Fig. S11C). Furthermore, anchorene largely rescued the loss of gravitropism caused by disruption of auxin transport in NPA treated seedlings (Fig. S11D and Fig. S11E). Using the *pPIN3::PIN3-GFP* marker line, we also examined the effect of anchorene on the auxin efflux carrier PIN3 that plays an important role in LR initiation and emergence (24). Application of anchorene led to a significant increase in PIN3 levels (Fig. 4E and 4F). However, the *pin3* mutant did not show altered ANR-RE (Fig. S11F), which may be due to redundancy of the PINs in regulating auxin transport.

### Application of anchorene leads to large scale changes in the collet transcriptome

Next, we performed RNA sequencing (RNA-Seq) on collets isolated from seedlings after treatment with anchorene, NPA, or RAM excision. NPA and anchorene treatment affected the expression level of 3355 overlapping genes and exerted the opposite effect on 2791 (83%) of them (Fig. 5A and 5C, Dataset S1). This is consistent with the opposite effects of NPA and anchorene on ANR development. In contrast, RAM excision and anchorene treatment led to a similar response in the expression level of 1459 genes and caused opposite effects in only 36 genes (Fig. 5B and 5D, Dataset S1), which is consistent with their common role in triggering ANR development. Biological Processes (BP) GO term analysis showed that many genes overlapping between anchorene treatment and RAM excision are related to auxin metabolism (Fig. S13, Dataset S2), indicating that modulation of this process is also important for regulating ANR development.

**Fig. 5.**
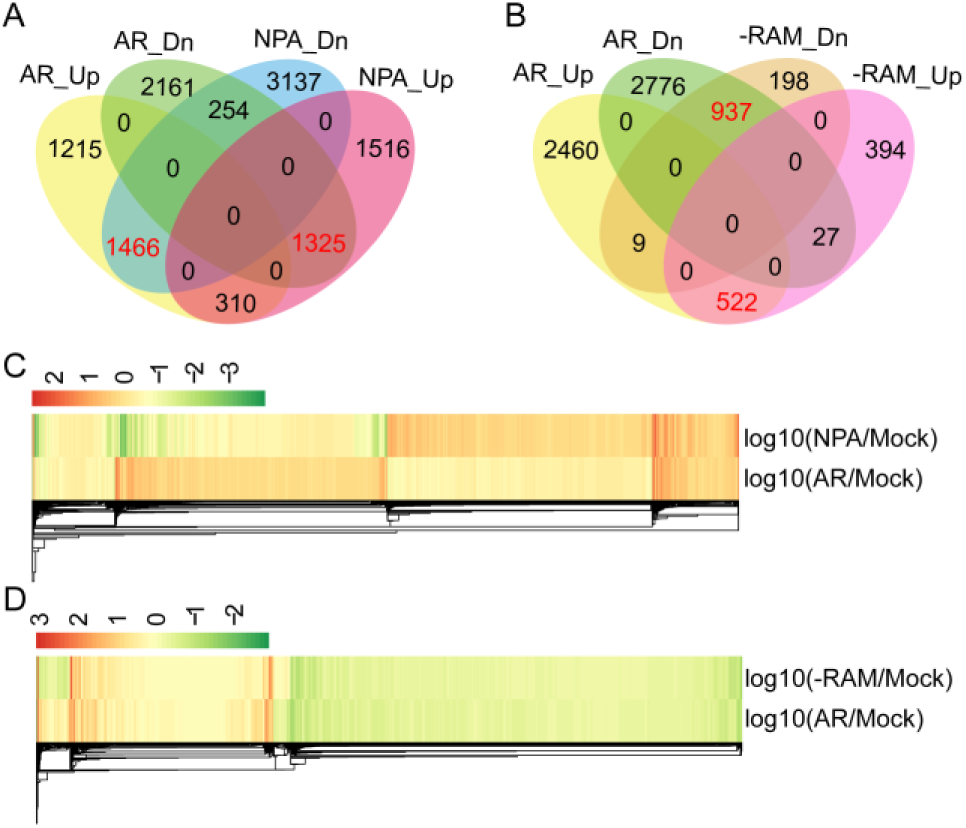
RNA-Seq data show that anchorene works antagonistically with NPA but mimics RAM excision (-RAM) at transcriptional level in the collet. The Venn diagrams show the numbers of the down-regulated (Dn) and up-regulated (Up) genes that overlap between anchorene treatment and NPA treatment (A), and -RAM (B). Heatmap clustering shows that the majority of anchorene and NPA overlapping genes are regulated in an oppositional expression pattern (C), while the majority of anchorene and -RAM overlapping genes are regulated in a similar pattern (D). AR: anchorene.

### Biological function of ANRs

To explore the biological role of ANRs, we tested the effect of soil type on ANR development. We compared ANR formation in seedlings grown in organic and sandy soil. Interestingly, about 63±17.5% of seedlings grown in sand formed ANRs at 8 dps, compared to only 4±3.5% of those grown in organic soil (Fig. 6A and 6B), suggesting that local environment is an important factor for ANR development. We analyzed the pH value, nutrient and element composition of the sand soil and organic soil, which showed quite different compositions (Table S1 and S2). Phosphorus and nitrogen are very important nutrient elements that regulate root development (42), so we compared the anchor root formation under phosphorus or nitrogen deficient conditions. Interestingly, nitrogen deficient but not phosphorus deficient condition showed elevated anchor root formation compared to mock condition. Then we further quantified the anchorene content under nitrogen deficient and mock conditions. Consistently, anchorene content was much higher under nitrogen deficient condition than that of mock, which indicates anchorene content is regulated by nitrogen status.

**Fig. 6.**
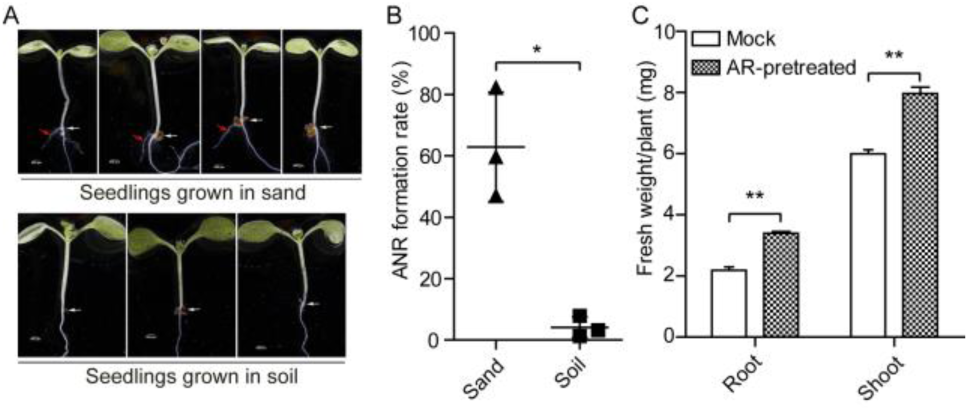
Biological functions of ANRs in Arabidopsis. (A) 8-day-old sand and soil grown seedlings are shown. White arrows indicate the collets, and the red arrows indicate the ANRs. (B) ANR formation rate of seedlings grown in sand and soil. (C) Anchorene pretreatment increased root and shoot biomass of Arabidopsis seedlings. Seedlings were grown in half MS agar plates with or without anchorene for 7 days, then seedlings were transferred to new plates without anchorene and grown for another 10 days. In B and C, data are presented as mean ± SD (two-tailed paired Student *t*-test, \**P* <0.05 \*\**P* <0.01) from three independent experiments. AR: anchorene.

Finally, we investigated the effect of anchorene treatment on plant growth by measuring root and shoot biomass in 17-day-old anchorene treated seedlings. A treatment with anchorene for one week led to increased ANR number, wider root system, and enhanced root (about 50%) and shoot (about 30%) fresh biomass (Fig. 6C and Fig. S15). These results indicate that anchorene can facilitate plant growth.

## Discussion

Arabidopsis forms three types of post-embryonic roots, ANRs, LRs and adventitious roots (18, 19). In this study, we examined the development of ANRs, the least characterized of these root types, and demonstrate that their formation is triggered by a carotenoid-derived signal. Moreover, we show that anchorene is the signal that regulates ANR development. To our knowledge, anchorene is the first reported diapocarotenoid with a specific regulatory function. Due to their instability and reactivity, diapocarotenoids have attracted little attention, and have mainly been studied as precursors of pigments such as crocetin (43). Hence, the identification of anchorene is expected to facilitate the discovery of further diapocarotenoid-based plant regulatory compounds and unravel new functions of carotenoid-derived metabolites.

Anchorene is a specific inducer of ANR formation, as shown by the inactivity of its isomer and derivatives, i.e. the corresponding dialcohol, diacid and diethyl ester (Fig. S3H and S3I). Unlike LRs, ANRs originate from the collet, which emerges from embryonic tissue. Therefore, ANR development is fundamentally different from that of LRs, which initiate near the root meristem (21). Although both ANRs and LRs originate from the pericycle, the collet pericycle forms only one or two ANRs located opposite each other, while the primary root pericycle continuously develops alternating LRs.

Anchorene is a synthetic compound that could arise *in planta* from carotenoid cleavage. Our work demonstrates that ANR development requires a carotenoid-derived signal (Fig. 2) and that anchorene exerts the function of this signal in carotenoid-deficient seedlings (Fig. 3A and 3B). Further, we show that anchorene is a natural Arabidopsis metabolite and its formation could be inhibited by NF treatment (Fig. 3C, Fig. S10D and S10E). The question of how anchorene is produced from carotenoids remains elusive. Theoretically, anchorene can arise by cleaving C11-C12 and C11’-C12’ double bonds in all carotenoids starting from *ζ*-carotene in the carotenoid biosynthesis pathway (Fig. S2). In contrast to ABA and SL, which derive from specific 9-*cis*-carotenoids (7, 8), anchorene can be formed from every *trans*- or *cis*-carotenoid with a continuously conjugated, *trans*-configured central moiety (C11 to C11’; see Fig. S2).

Several CCDs from plants (16), fungi (44) and cyanobacteria (17) produce diapocarotenoids, either by repeated cleavage of carotenoid substrates or by specifically targeting apocarotenoids. However, enzymatic studies on Arabidopsis CCDs do not support the formation of anchorene by a single CCD. Indeed, none of the Arabidopsis CCDs are capable of performing cleavage at both of the C11-C12 and C11’-C12’ double bonds *in vitro* (45), but interestingly, our results showed that OH-APO12’ is likely a specific precursor of anchorene in Arabidopsis by feeding experiments (Fig. S12). It was previously shown that OH-APO12’ could be produced by a maize NCED enzyme, VP14, by the cleavage of zeaxanthin (7), however individual Arabidopsis *ccd* or *nced* mutants did not show ANR-RE reduction (Fig. S7B and S7F). Arabidopsis has five NCEDs, which make it difficult to knock out all of these NCEDs to test their involvement in anchorene production, so we cannot exclude that anchorene is redundantly produced by more than one CCD or CCD combination that exerts the required cleavage activity *in planta*. Carotenoids are also cleaved non-enzymatically, catalyzed by ROS (15). To determine whether anchorene can be produced non-enzymatically, we tested whether it was formed as oxidation product of synthetic β-carotene in an organic solution *in vitro*. LC-MS analysis identified an anchorene pattern similar to that observed in shoot samples (Fig. S16), indicating that anchorene may also be produced non-enzymatically *in planta*. A number of apocarotenoid signaling molecules have been shown to be produced non-enzymatically *in vitro*, both in plants and animals (17, 46). This may indicate that the specificity of anchorene activity is regulated mainly by a receptor, rather than biosynthetically.

Our study reveals a central role of auxin in ANR development. The auxin responsive transcription factors *ARF7* and *ARF19*, which are key regulators of LR initiation (26, 28), are also indispensable for ANR initiation (Fig. 4A and 4B). Moreover, the auxin analog NAA strongly increases ANR-RE while the auxin transport inhibitor NPA impedes the formation of ANRs (Fig. 4C), suggesting that auxin content and distribution are both important for ANR-RE. Using auxin-responsive marker lines, we showed that the application of anchorene increased the levels of auxin reporters both in ANR primordia and the surrounding tissue (Fig. S4 and Fig. S9), suggesting that anchorene triggers ANR formation by modulating auxin transport and levels. Consistent with these results, transcriptome analysis showed that anchorene application and removal of RAM lead to the induction of many auxin biosynthesis genes (Fig. S13). Future work will shed light on anchorene’s mode of action. However, it can be speculated that anchorene may act via post-translational modification of regulatory modulate proteins by building conjugates with lysine and/or cysteine residues, as proposed for other carotenoid cleavage products (47).

Root systems are important not only for anchoring plants in soil but also for absorbing water and nutrients. We observed an increase in ANR number upon using nutrient-poor sandy soil (Fig. 6A and 6B). We additionally characterized a growth-promoting effect of anchorene (Fig. 6C). Moreover, we examined more anchorene and anchor root formation under nitrogen deficient conditions compared to those of normal growing seedlings, indicating that ANRs may improve nutrient uptake and pointing to potential for applications in agriculture or horticulture. It is worth mentioning that the carotenoid cleavage product anchorene is a commercially available compound used as building block for the manufacturing of different carotenoids on an industrial scale (48).

## Materials and Methods

Experimental procedures for chemical synthesis and preparation, plant materials and growth conditions, root phenotyping assays, and other experiments performed in this study are described in SI Appendix, Supplementary Materials and Methods.

## Supporting information

## Acknowledgements

We thank Dr. Ikram Blilou and Dr. Bernd Schaefer for valuable discussions, and Mohammed Khalid for technical support. This work was supported by Baseline funding and the Research Grants Program-Round 4 (CRG4)) from King Abdullah University of Science and Technology (KAUST) to S.A., by the Arnold and Mabel Beckman Postdoctoral Fellowship (AJD) and by the Howard Hughes Medical Institute and the Gordon and Betty Moore Foundation (through Grant GBMF3405) to PNB.

## Author contributions

K.J., A.J.D., P.N.B. and S.A. conceived the study; J.M., N.M.K. and X.G. performed anchorene LC-MS analysis; G.C and M.A. performed the RNA-Seq analysis; E.S. and M.R. synthesized the anchorene derivatives; K.J. and A.J.D. performed the other experiments and analyzed the data; K.J., A.J.D., P.N.B. and S.A. wrote the paper.

