## Supplementary material for "Anchorene is an endogenous diapocarotenoid required for anchor root formation in Arabidopsis"

**This PDF file includes:**

Supplementary text
Figs. S1 to S16
Tables S1 to S2
References for SI reference citations

### Supplementary text

#### Materials and Methods

##### Chemicals

Diapo1, Diapo2, Diapo4, Diapo5 and Diapo6 (Supplemental Figure 1) were synthesized by Buchem (Netherlands). Diapo3 (anchorene) was firstly custom synthesized by Buchem (Netherlands), and was later synthesized together with D6-anchorene and anchorene derivatives according to the protocol (s. below). Apocarotenoids and diapocarotenoids were dissolved in acetone to make 10 mM stock solutions. For the screening experiments, each chemical was diluted in half strength Murashige and Skoog (MS) media (with 0.5 % sucrose + 1% agar, 0.5 g/L MES, pH 5.7) to reach indicated concentrations of 25 μM and 5 μM. Stock solutions for D15 (100 mM), Norflurazone (NF, 10 mM) (Chem Service), and 2-(4-chlorophenylthio)-triethylamine hydrochloride (CPTA, 50 mM) [obtained from the laboratory of L. E. Sieburth at the University of Utah (Salt Lake City, UT)] were prepared in dimethyl sulfoxide (DMSO). 125 μM D15, 1μM NF, and 100 μM CPTA working solutions were obtained by diluting the corresponding stock solutions in a fore mentioned MS medium. GR24 (Chiralix, Netherlands) was prepared in acetone. Abscisic acid (ABA), naphthaleneacetic acid (NAA) and 1-N-naphthylphthalamic acid (NPA) were purchased from Sigma Aldrich, and their stock solutions were all prepared in water at 1 mM concentration.

##### Synthesis of anchorene, D6-anchorene and anchorene derivatives

Unless otherwise noted, all commercially available compounds were used as provided without further purification. $CH_2Cl_2$ and THF used for the reactions were purified by an MBraun solvent purification system (SPS). Solvents for chromatography were technical grade and freshly distilled prior to use. Analytical thin-layer chromatography (TLC) was performed on Merck silica gel aluminium plates with F-254 indicator, visualized by irradiation with UV light. Column chromatography was performed using silica gel (Macherey Nagel, particle size 0.040-0.063 mm). Solvent mixtures are understood as volume/volume. 1H-NMR and 13C-NMR were recorded on a Varian AV400 or AV600 spectrometer. Data are reported in the following order: chemical shift (δ) in ppm and coupling constants (J) are in Hertz (Hz). IR spectra were recorded on a Perkin Elmer-100 spectrometer and are reported in terms of frequency of absorption (cm-1). Mass spectra (EI-MS, 70 eV) were conducted on a Finnigan SSQ 7000 spectrometer.

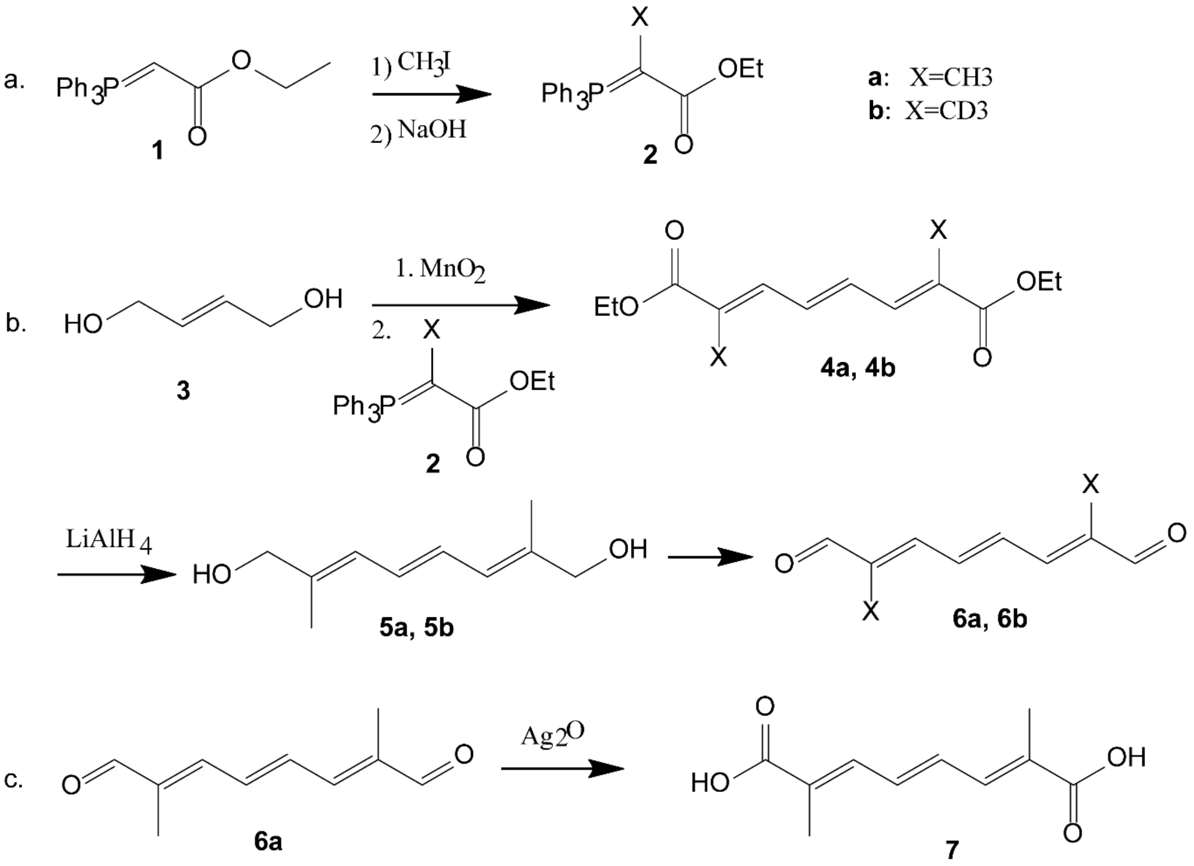


Supporting Fig. 1: The synthetic route for anchorene, D6-anchorene and anchorene derivatives.

**4a, 4b** (AR diethyl ester): (2E, 4E, 6E)-diethyl-2,7-dimethylocta-2,4,6-trienedioate (1, 2)

But-2-ene-1,4-diol 1 (1 equiv) dissolved in $CH_2Cl_2$ was added to a solution of $MnO_2$ (18 equiv) in $CH_2Cl_2$ at 0 °C. Phosphorane 2 (2.4 equiv) dissolved in DCM was then added. The reaction mixture was stirred at room temperature until the TLC showed the full consumption of the starting material. $MnO_2$ was removed by filtration over celite and the filtrate concentrated in *vacuo*. Purification by column chromatography (hexane: EtOAc 5:1) gave the product 3 as white crystalline solid.

**4a**: 1H NMR (400 MHz, CDCl3): δ (ppm): 1.31 (t, J = 7.2 Hz, 6 H), 2.0 (d, J = 7.2 Hz, 6 H), 4.22 (q, J = 7.2 Hz, 4H), 6.79 (dd, J = 7.6 Hz/2.8 Hz, 2 H), 7.28 (dd, J = 8.0 Hz/1.6 Hz, 2H).

**4b**: 1H NMR (600 MHz, CDCl3): δ (ppm): 1.32 (t, J = 7.2 Hz, 6 H), 4.23 (q, J = 7.2 Hz, 4H), 6.79 (dd, J = 7.8 Hz/3.0 Hz, 2 H), 7.29 (dd, J = 7.8 Hz/3.0 Hz, 2H). 13C NMR (150.9 MHz, CDCl3): δ (ppm) :14.3, 60.8, 130.1, 133.6, 137.2, 168.0. MS (EI) m/z (%): 258.2 [M+.] (100). IR (ATR): ṽ = 2114, 1694, 1615, 1475, 1368, 1284, 1225, 1101, 995 $cm^{-1}$.

**5a** (AR dialcohol): (2E, 4E, 6E)-2,7-dimethylocta-2,4,6-triene-1,8-diol

Ester 3 (1 equiv) in dry THF was added to a suspension of $LiAlH_4$ (2.4 equiv) in dry THF at 0 °C. The reaction mixture was stirred at this temperature for 1 h. The reaction was quenched by slow addition of water and 20% NaOH solution. The organic phase was separated, and the aqueous phase washed with EtOAc. The combined organic phase was dried over $MgSO_4$ and concentrated leading to the desired diol 4.

**5a**: 1H NMR (600 MHz, CDCl3): δ (ppm) = 1,61 (s, 2H), 1.81 (s, 6H), 4,10 (s, 4H), 6.16 (d, J = 7.8 Hz, 2H), 6.45 (dd, J = 7.2 Hz/3.0 Hz, 2 H).

**6a, 6b** (anchorene) (2E,4E,6E)-2,7-dimethylocta-2,4,6-trienedial (3)

To the suspension of $LiAlH_4$ (2.4 equiv) in dry THF at 0 °C was added the ester **4** (1 equiv) in dry THF. The reaction mixture was stirred at this temperature for 1 h. The reaction was quenched by slow addition of water and 20% NaOH solution. The organic phase was separated and the aqueous phase washed with EtOAc. The combined organic phase dried over $MgSO_4$ and concentrated. The residue **5** was oxidized without further purification. To a cooled solution of the crude diol **5** in acetone was added $MnO_2$ (18 equiv). The reaction mixture was allowed to warm up to room temperature and stirred for 24 h. The solid was removed by filtration over a pad of celite and washed with $CH_2Cl_2$. The solvent was removed in vacuo and the residue was purified by column chromatography ($SiO_2$, hexane:EtOAc 5:1). The dialdehyde **6** was isolated as yellow solid.

**6a**: 1H NMR (600 MHz, $CDCl_3$): δ (ppm) = 1.93 (s, 6H), 6.96 - 7.02 (m, 2H), 7.07 (dd, J = 7.8 Hz/ 3.0 Hz, 2H), 9.53 (s, 2 H).

**6b**: 1H NMR (400 MHz, CDCl3): δ (ppm) = 6.99 (dd, J = 8.4 Hz/ 2.8 Hz,2H), 7.07 (dd, J = 8.0 Hz/ 3.2 Hz, 2H), 9.55 (s, 2 H). 13C NMR (100.5 MHz, CDCl3): δ (ppm) = 134.4, 140.9, 146.0, 194, 4. MS (EI) m/z (%) = 170.1 [M+.] (100). IR (ATR): ṽ = 2078, 1718, 1662, 1369, 1270, 1171, 1032, 979 $cm^{-1}$.

**7** (AR diacid): (2E, 4E, 6E)-2,7-dimethylocta-2,4,6-trienedioic acid (4)

A mixture of dial **6a** (6 mmol, 1 equiv), $Ag_2O$ (9.1 mmol, 1.4 equiv), 3 mL NaOH (10% solution) and 20 mL $H_2O$ was refluxed for 24h. The reaction mixture was diluted with 50 mL $H_2O$, and the solid was separated by filtration. The filtrate was acidified with diluted $HNO_3$ and the resulting solid filtered up and recrystallized from ethanol to give the product as off-white solid. 1H NMR (600 MHz, DMSO-d6): δ (ppm) = 1.92 (s, 6H), 7.12 - 7.32 (m, 4H), 12.4 (bs, 2H).

#### Plant materials and growth conditions.

Col-0 was used as wild type, unless otherwise noted. Mutants *psy, isph1, lut1, lut2, ccd1, ccd4, ccd7, ccd8, nced2, nced3, nced5, nced6, nced9, aba1, aba3* were described previously (5). The mutant *pin3-4* were acquired from ABRC stock center. The *arf7arf19* (CS24625) mutants were acquired from the European Arabidopsis Stock Centre and as descripted previously (6). *pWOX5::GFP* (7), *pDR5::LUC* (8) *pDR5rev::GFP* (9) and *pPIN3::PIN3-GFP* (10) transgenic marker lines were all as described previously.

Supporting table 1: Mutants and marker lines used in this study.

| Alleles name | Gene locus | Description | References |
|---|---|---|---|
| *psy-1* (Salk_054288) | AT5g17230 | Knock out mutant | 5 |
| *ispH-1* | AT4g34350 | Knock out mutant | 5 |
| *lut1* | AT3G53130 | Knock out mutant | 5 |
| *lut2* | AT5G57030 | Knock out mutant | 5 |
| *ccd1-1* | AT3g63520 | Knock out mutant | 5 |
| *ccd4-1* | AT4g19170 | Knock out mutant | 5 |
| *ccd7 (max3-11)* | AT2g44990 | Knock out mutant | 5 |
| *ccd8 (max4-6)* | AT4g32810 | Knock out mutant | 5 |
| *nced2-3* (Salk_090937) | AT4g18350 | Knock out mutant | 5 |
| *nced3* (N3KO-6620) | AT3g14440 | Knock out mutant | 5 |
| *nced5* (N5KO-4250) | AT1g30100 | Knock out mutant | 5 |
| *nced6* (WISC.DSLox471G6) | AT3g24220 | Knock out mutant | 5 |
| *nced9* (Salk_051969) | AT1g78390 | Knock out mutant | 5 |
| *aba1-6* (CS3772) | AT5g67030 | Knock out mutant | 5 |
| *aba3-1* (CS157) | AT1g16540 | Knock out mutant | 5 |
| *pin3-4* (SALK_038609) | AT1G70940 | Knock out mutant | Identified by SALK |
| *arf7arf19* | AT5G20730 & AT1G19220 | Knock out mutant | 6 |
| *pWOX5::GFP* | - | *WOX5* promoter marker line | 7 |
| *pDR5::LUC* | - | synthetic auxin marker line | 8 |
| *pDR5Rev::GFP* | AT3G11260 | synthetic auxin marker line | 9 |
| *pPIN3::PIN3-GFP* | AT1G70940 | *PIN3* marker line | 10 |

Sterilized Col-0 and mutants seeds were kept at 4°C in darkness for 3 days to stimulate seed germination and then sown on half strength MS (with 0.5 % sucrose + 1% agar, 0.5 g/L MES, pH 5.7) plates supplemented with the indicated compounds. Plates were vertically grown in Percival growth chambers under long day (16 h light/8 h dark, 22°C, 60% relative humidity, light density: 4000 LUX) LED white light [Hyperikon 16W LED Light Bulb A21, 16W (100W Equivalent), CRI92, 1620 Lumens, 4000K (Daylight Glow)] conditions. Light fluorescence rates were measured using a digital LUX meter (PeakTech 5025).

##### ANR, lateral root and primary root phenotyping assays

ANRs were counted in seedlings vertically grown on half strength MS (with 0.5 % sucrose + 1% agar, 0.5 g/L MES, pH 5.7) media 8 days post-stratification (dps) using a dissection microscope. To investigate ANR formation after root apical meristems (RAM) excision (ANR-RE), RAM of seedlings were excised using sterile scalpels at 5 dps and then grown for another 3 days prior to counting the number of emerged ANRs. Protocol for quantifying LR capacity has been described previously (5). Specifically, RAM of seedlings were excised at 8 dps and then grown for another 3 days prior to counting the number of emerged LRs. For determining the effect of anchorene on lateral roots, the number of emerged lateral roots were counted using a dissection microscope at 8 dps. Primary roots length was measured using the publicly available ImageJ software (http://rsbweb.nih.gov/ij/) after taking digital photographs.

For the quantification of ANR formation under OH-Apo10’ and OH-Apo12’ treatment conditions, the sterilized

Arabidopsis seeds were exposed to light for 24 hours first, and then plated to 1/2MS plates with indicated chemicals; keep the plates under darkness (22$^{o}$C) for another two days and then exposed to long day light conditions for another 7 days to count the ANR emergence.

##### Confocal microscopy

To examine ANR initiation and primordium, the ClearSee protocol was applied as described previously (11). Briefly, seedlings exposed to various treatments were fixed using 4% paraformaldehyde dissolved in phosphate buffered-saline (PBS) for 30 minutes. Fixed seedlings were washed twice with PBS and then immersed in ClearSee solution (10% w/v Xylitol, 15% w/v sodium deoxycholate, and 25% w/v urea in water). After incubating the seedlings in ClearSee solution in the dark at room temperature for 2-3 days, laser scanning confocal microscopy (Zeiss LSM 510 microscope) was used to examine the roots. To examine *pDR5*$_{Rev}$*::GFP* fluorescence at the collet region, live seedlings were directly used for laser scanning confocal microscopy (Zeiss LSM 510 microscope) examination.

For fluorescence intensity quantification of *pDR5*$_{Rev}$*::GFP* and *pPIN3::PIN3-GFP* marker lines, ImageJ software (http://rsbweb.nih.gov/ij/) was used after taking confocal microscopy photos. All calculations are background subtracted.

##### Luciferase assay

Luciferase activity was assayed as previously described (4). Briefly, 1 mL of 5 mM Potassium Luciferin (Gold Biotechnology) dissolved in water was directly applied to *pDR5::LUC* seedlings grown vertically on half strength MS plates. The luciferin solution was allowed to dry for 5-10 minutes in the dark, at room temperature. Seedlings were then imaged using a Lumizone CA automated Chemiluminescence system. Seven minute exposure times were used.

##### Qualitative and quantitative identification of anchorene using LC-MS

30-40 mg of freeze-dried and ground Arabidopsis seedlings tissues were extracted using 1 mL of acetonitrile with anti-oxidant [0.1% butylated hydroxytoluene (BHT)] for 15 min in an ultrasonic bath (Branson 5510EDTH, 25 $^{o}$C), followed by centrifuge for 8 min at 13000 rpm at 4 °C. The collected supernatant was dried using a concentrator (Labconco RapidVap System). The extract was derivatized (see the following Chemical reaction equation, Supporting Fig. 2) according to the protocol described previously with minor modification (12). 50 µL of derivatization solution containing 5 mg/mL derivatization reagent ($N^2$,$N^2$,$N^4$,$N^4$-tetraethyl-6-hydrazineyl-1,3,5-triazine-2,4-diamine) (Chemspace) and 1% formic acid in methanol was kept at 37 °C for 15 min. Then the sample solution was diluted to 150 µL with 1% formic acid in methanol and filtered by 0.22 µm filter before LC-MS analysis.

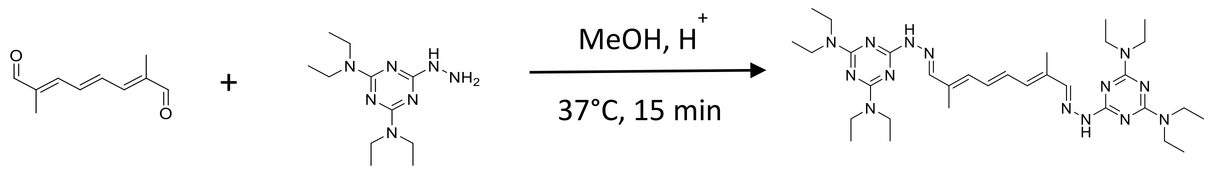


Supporting Fig. 2: The derivatization reaction for anchorene.

The qualitative analysis of derivative anchorene were performed on UHPLC-Q-Exactive Plus MS. Chromatographic separation was achieved on an Acquity UPLC BEH $C_{18}$ column (100 × 2.1 mm; 1.7 µm; Waters) using a mobile phase consisting of water:acetonitrile (95:5, v:v, A) and pure acetonitrile (B), both containing 0.1% formic acid. A gradient was applied, starting with 40% B and increasing to 50% B over 15 min. A concentration of 50% B was maintained for 10 min and then was increased to 100% B over 4 min and then maintained for 5 min. To equilibrate the column before the next run, the mobile phase was adjusted back to 40% B in 1 min and this concentration was maintained for 5 min prior to the next sample injection. A flow rate of 0.2 mL/min and a column temperature of 35 °C were maintained throughout the run. The eluent of the column was introduced to the mass spectrometer using heated-electrospray ionization in positive mode. The injection volume was 15 µL. The conditions of the mass spectrometer were set as following: resolution: 280,000; automatic gain control: $3\times10^6$; maximum injection time: 200 ms; sheath gas flow: 40 arbitrary units; auxillary gas flow: 10 arbitrary units; spray voltage, 4.0 kV; capillary temperature: 300 °C; vaporizer temperature: 300 °C. The quantitative analysis of derivative anchorene were carried out on HPLC-Q-Trap MS/MS. Chromatographic separation was achieved on an Acquity UPLC CSH $C_{18}$ column (50 × 2.1 mm; 1.7 µm; Waters) using a mobile phase consisting of water:acetonitrile (95:5, v:v, A) and pure acetonitrile (B), both containing 0.1% formic acid. A gradient was applied, starting with 10% B and increasing to 40% B over 5 min. Then 15% B was increased within 10 min followed by an increase of 45%B over 2 min. And then 100% B was maintained for 10 min. To equilibrate the column before the next run, the mobile phase was adjusted back to 10% B in 1 min and this concentration was maintained for 8 min prior to the next sample injection. A flow rate of 0.15 mL/min and a column temperature of 40 °C were maintained throughout the run. The eluent of the column was introduced to the mass spectrometer using turbo spray ion source in positive mode. The injection volume was 10 µL. The conditions of the mass spectrometer were set as following: curtain gas: 30; ionspray voltage: 5 kV; temperature: 400 °C; ion source gas 1: 30; ion source gas 2: 40; declustering potential, 55; entrance potential: 10; collision energy: 25; collision cell exit potential: 10. For derivative anchorene: Q1 mass (Da): 635.5, Q3 mass (Da): 239.2; for derivative $D_6$-anchorene: Q1 mass (Da): 641.5, Q3 mass (Da): 239.2.

##### RNA-Seq materials preparation and data analysis

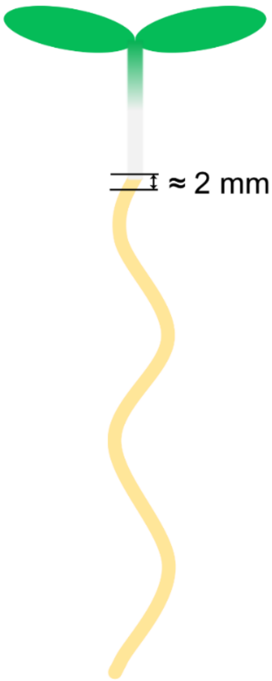


Supporting Fig. 3: About 2 mm collet region was excised for RNA-Seq sampling.

For anchorene and NPA treatment, Col-0 seedlings were vertically grown on half MS (with 0.5% sucrose + 1% agar, 0.5 g/L MES, pH 5.7) plates supplemented with 20 μM anchorene or 1 μM NPA for 5 days. RAM excision treatment was applied by excising the RAM from 4.5 dps Col-0 seedlings, and samples (2 mm long sections of at the collet region, see Supporting Fig. 3) were collected 12 hours after excision. Sections from approximately 50 seedlings were used for each RNA sample. Total RNA was extracted using Direct-zol RNA MiniPrep Plus (200 Preps) w/ Zymo-Spin (ZYMO research). 100 ng total RNA was used for RNA Hi-seq 4000. For each treatment, we collected 3 RNA samples isolated from 3 independent experiments.

Prior to the analysis of RNA-Seq data, the adaptor sequences and low-quality ends of sequenced reads were trimmed using Trimmomatic v0.32 (13) and quality-checked using FastQC v0.11.3 (14). To quantify the expression level of genes, the remaining reads were pseudo-aligned to the publicly available TAIR10 *A. thaliana* transcriptome (release 34) using kallisto v0.43.0 (15). The estimated read counts and calculated transcripts per million (TPM) were subsequently passed to sleuth v0.28.1 (16) for differential expression analysis. Significantly differentially expressed genes were identified based on a cutoff of fold-change > 1.5 and q-value < 0.05. Gene ontology and enrichment analysis was conducted using clusterProfiler (17). RNA-Seq data can be accessed at NCBI via BioProject ID PRJNA489360.

##### ANR phenotyping in sand and soil

Silver sand (VWR) and soil (Asdcofert.com) were used for ANR phenotyping experiments. Col-0 seeds were kept in 4 °C for 3 days to insure uniform germination, and then sown in pots with sand or soil. The pots were kept under long-day photoperiod conditions (16 h light/8 h darkness, 22ºC, 60% humidity) for 8 days. Then, forceps were used to gently take the seedlings from the sand or soil. Seedlings were then washed by water, prior to counting the ANRs under microscope.

##### Nutrient elements and pH value analysis in sand and soil

For micro nutrient elements analysis: ICP-OES was used to measure elemental levels of Fe, K, Mg, Mn, P and Zn. Specifically, homogenized dry soil (50-100 mg) and sand (100-200 mg) samples were dissolved in the vessels with 8 ml nitric acid together with negative control which only include nitric acid and samples spike with all the standard (6 ml nitric acid + 2 ml standards mixture). Samples were digested using Microwave digestion. After the digestion, $ddH_2O$ was added for a final volume of 25 mL. Samples were centrifugated and the supernatant was collected for ICP-OES analysis.

For the C, H, N, S elemental analysis, 5-20 mg homogenized dry soil and sand samples were analyzed by CHNS and Sulfanilmide was used as a standard.

For pH measurements, 15 ml soil or sand were soaked in 30 ml PH7.0 ddH2O. The samples were vortexed and incubated in root temperature overnight. 15 ml supernatant from each sample was used to measure pH values by a standard pH meter.

##### Hoagland and nutrient deficient growing conditions

Col-0 seedlings were grown in vertical Hoagland medium (with 0.5 % sucrose + 1% agar, 0.5 g/L MES, pH 5.7) plates for 10 or 12 dps. For phosphorus deficient medium, remove the $KH_2PO_4$ and add equal content of KCl in Hoagland solution; for nitrogen deficient medium, remove the $NH_4NO_3$ in Hoagland solution.

##### Determination of Biomass

Col-0 seedlings were grown in vertical half MS (with 0.5 % sucrose + 1% agar, 0.5 g/L MES, pH 5.7) plates with or without anchorene application (20 μM) for one week. Selected well-growing mock seedlings (without anchorene treatment) and anchorene treated seedlings with two emerged ANRs were then transferred to new half MS media plates, and 12 seedlings per plate were grown vertically for another 10 days. After 10 days, roots and shoots were separately collected as one technical replicate from each plate. Four technical replicates for each treatment were quantified in each experiment and three independent experiments were conducted.

##### RNA extraction and Quantitative Real Time PCR

Total RNA was extracted from 10-20 seedlings grown under indicated conditions, using Direct-zol RNA MiniPrep Kit (Zymo Research). 1 µg total RNA was reverse-transcribed using iScript™ Reverse Transcription Supermix for RT-qPCR kit (Bio Rad). Amplification was carried out with SYBR® Green Real-Time PCR Master

Mixes kit (Life technologies). Quantitative Real time PCR was performed in a StepOne™ Real-Time PCR Systems (Life Technologies). The thermal profile for real-time PCR was 95°C for 2 min, followed by 40 cycles of 95°C for 15s and 60°C for 30s. Carotenoids biosynthesis genes *PSY*, *PDS*, *LYC*, *ZDS*, and *CCR2* were amplified. *CACS* was used as reference gene. Primers used for these genes amplification are listed in Table S3.

#### Carotenoids extraction and quantification

About 10 mg of freeze-dried and ground Arabidopsis seedlings with or without anchorene treatment were extracted using 2 mL of acetone with anti-oxidant (0.1% BHT) for 20 min in an ultrasonic bath (Branson 5510EDTH, 25 $^{o}$C), followed by centrifugation for 8 min at 13000 rpm at 4 °C. The collected supernatant was diluted 10 times in acetone for Spectrophotometer analysis. The quantification of total carotenoids by absorption spectrum was referred to previous described protocol (18). Specifically, absorption wavelength at 661.6 nm, 644.8 nm and 470 nm were measured. The total carotenoids was calculated according to the formula: $C_a$=11.24$A_{661.6}$-2.04$A_{644.8}$; $C_b$=20.13$A_{644.8}$-4.19$A_{661.6}$; total carotenoids (μg/ml) = (1000$A_{470}$-1.90$C_a$-63.14$C_b$)/214.

**Supplemental Figures:**

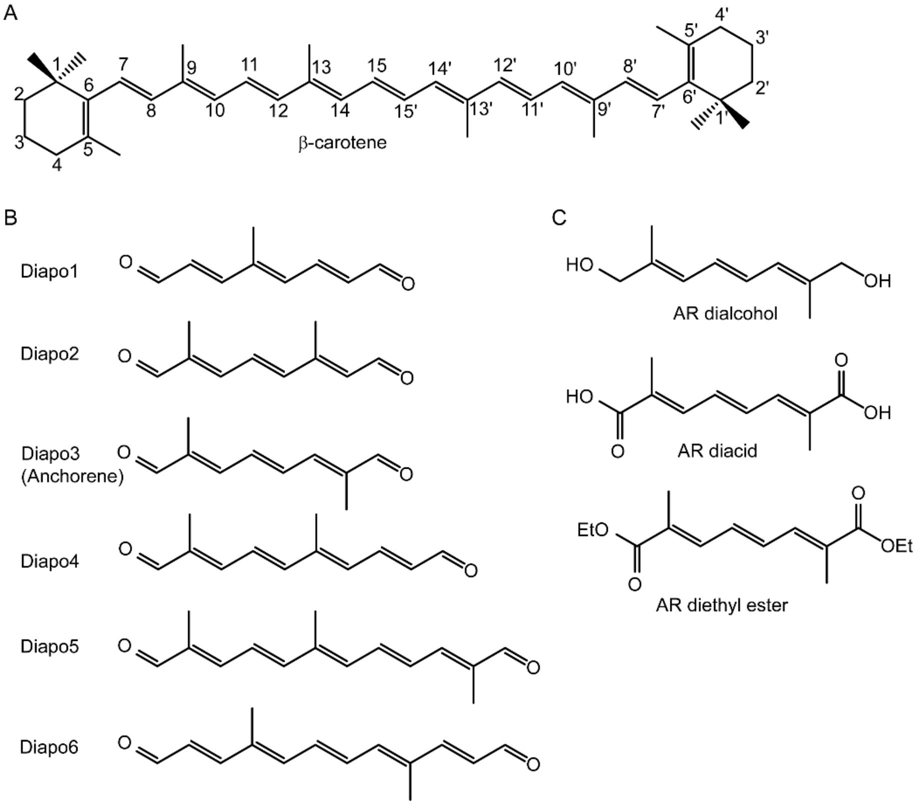


Fig. S1. Structures of carotenoid-derivatives. (A) β-carotene was used for the C-atom numbering. (B) structures of diapocarotenoids used in this study: Diapo1 [10,14’-diapocarotene-10,14’-dial; (2E,4E,6E)-4-methylocta-2,4,6-trienedial]; Diapo2 [8,15-diapocarotene-8,15-dial; (2E,4E,6E)-2,6-dimethylocta-2,4,6-trienedial]; Diapo3 [12,12’-diapocarotene-12,12’-dial; (2E,4E,6E)-2,7-dimethylocta-2,4,6-trienedial]; Diapo4 [8,14’-diapocarotene-8,14’-dial; (2E,4E,6E,8E)-2,6-dimethyldeca-2,4,6,8-tetraenedial]; Diapo5 [8,12’-diapocarotene-8,12’-dial; (2E,4E,6E,8E,10E)-2,6,11-trimethyldodeca-2,4,6,8,10-pentaenedial]; Diapo6 [10,10’-diapocarotene-10,10’-dial; (2E,4E,6E,8E,10E)-4,9-dimethyldodeca-2,4,6,8,10-pentaenedial]. (C) Structures of anchorene derivatives: AR dialcohol [(2E,4E,6E)-2,7-dimethylocta-2,4,6-triene-1,8-diol]; AR diacid [(2E,4E,6E)-2,7-dimethylocta-2,4,6-trienedioic acid]; AR diethyl ester [diethyl (2E,4E,6E)-2,7-dimethylocta-2,4,6-trienedioate].

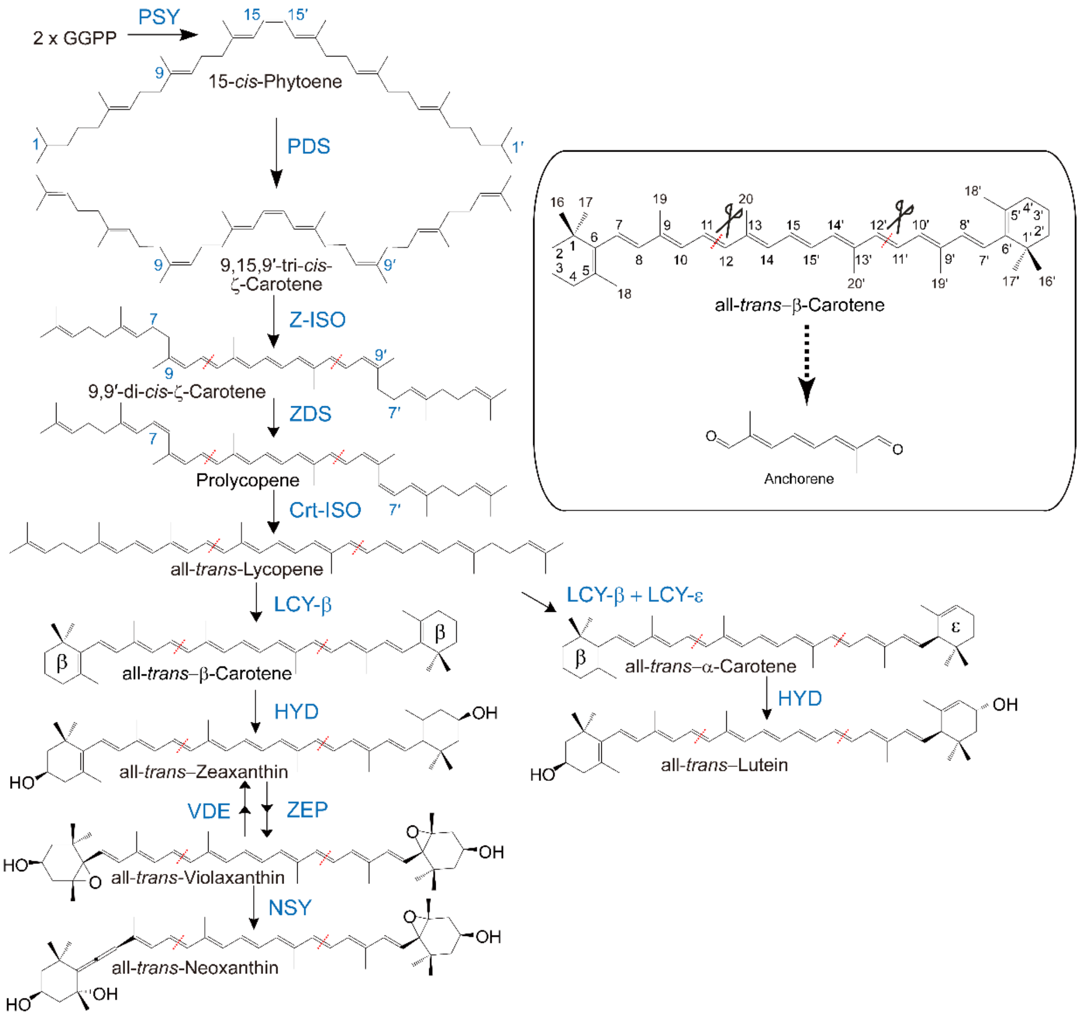


Fig. S2. Biosynthesis pathway of carotenoids and supposed precursors for anchorene in plants. (Inset) Proposed production of anchorene (12,12’-diapocarotene-12,12’-dial) from β-carotene by oxidative cleavage; all-trans-β-carotene is taken as an example for the C-atom numbering of carotenoids and the production of anchorene. Enzyme names are shown in blue; red dashed lines in carotenoids indicate the position that could be oxidatively cleaved to produce anchorene.

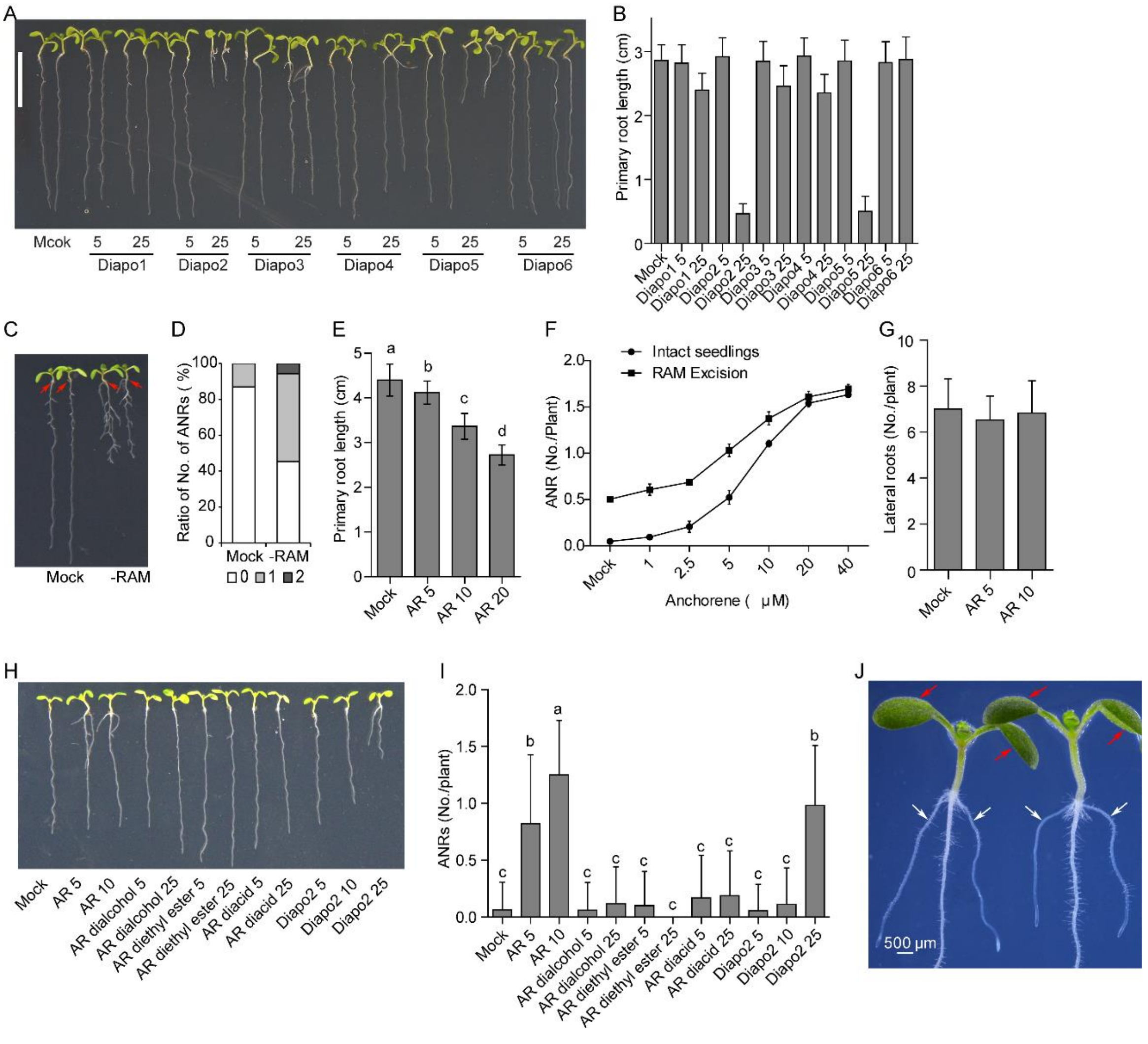


Fig. S3. Characterization of effects of putative carotenoid-derivatives on root development in Arabidopsis. (A) Representative Col-0 seedlings treated by different carotenoid-derivatives. Red arrows indicate emerged anchor roots (ANRs). (B) Quantification of primary root length of Col-0 seedlings treated by various diapocarotenoids. n=26, 26, 27, 25, 12, 26, 24, 28, 21, 28, 15, 28, 22 respectively. In A and B, 7 dps seedlings grown in half MS agar plates supplied with 5 µM or 25 µM of the indicated chemicals. Mock treatment consisted of acetone. (C) Representative Col-0 seedlings showed that RAM excision promotes ANR formation. (D) Quantification of ANRs formation with or without RAM excision as percentage of seedlings with 0, 1 and 2 ANRs formation. n=61, 53 respectively. (E) Anchorene has a minor inhibition on primary root length. 8 dps Col-0 seedlings were analyzed. (F) The dose response of anchorene effect on ANR formation under normal and RAM excision conditions. (G) Anchorene has no effect on lateral root formation. n=45 for each treatment. 8 dps seedlings were used for analysis. (H) Representative seedlings treated by different anchorene analog and derivatives. (I) Quantification of different anchorene analog and derivatives effects on ANR formation. Concentrations (µM)

for each chemical used in E-I are as indicated. (J) ANRs are always in the same geometric plane as cotyledons. White arrows indicate ANRs, and red arrows indicate cotyledons. In G and I, data were presented as mean ± SD (one-way ANOVA with Tukey multiple-comparison test, $P < 0.05$); different letters denote significant differences; in E, n=19, 20, 22, 23, respectively, and in I, n=48, 60, 52, 49, 51, 52, 48, 48, 54, 53, 55, respectively. Scale bar, 1cm.

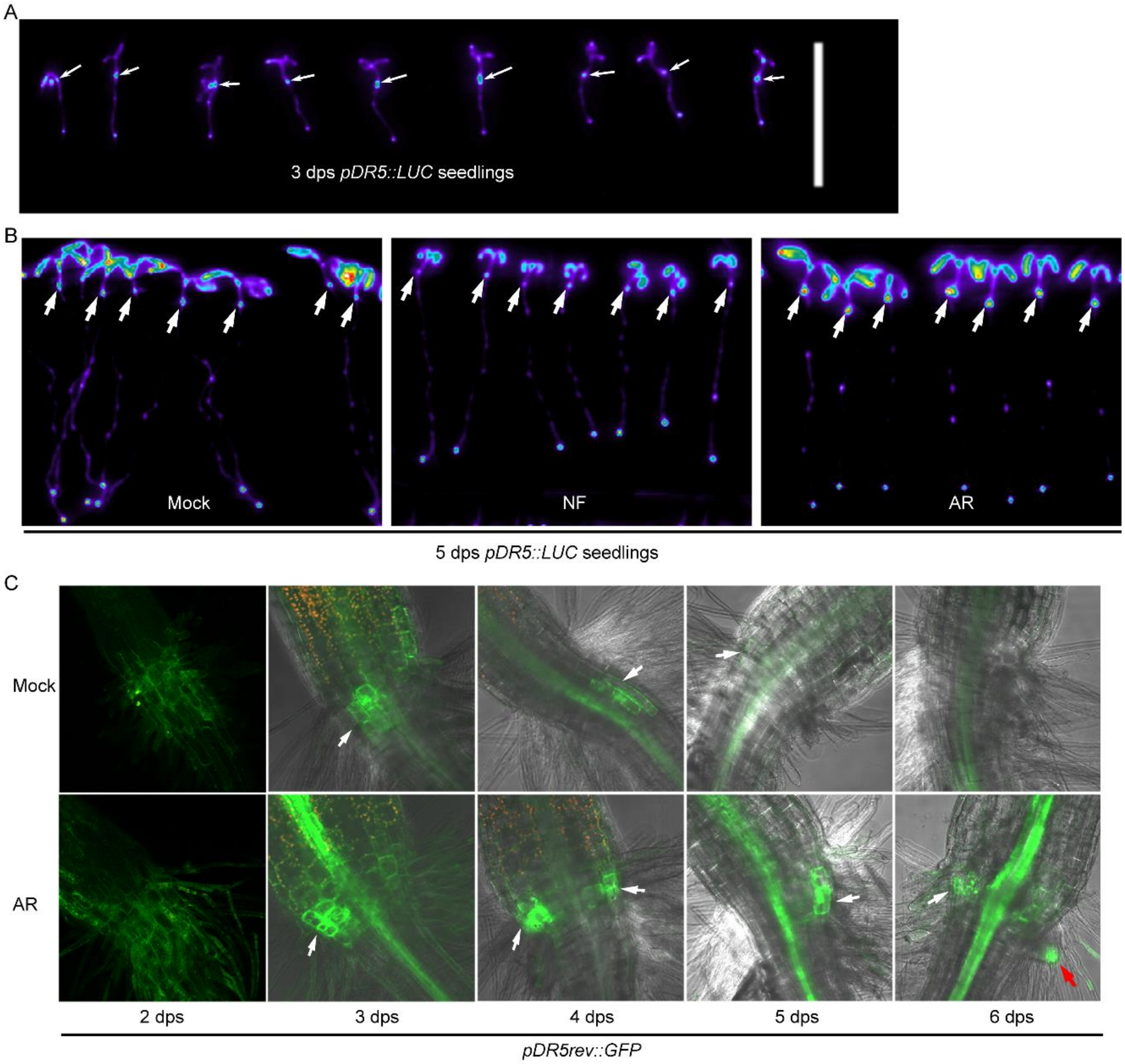


Fig. S4. Characterization of ANR development by different DR5 marker lines. (A) 3 dps *pDR5::LUC* seedlings indicate the ANR initiation. (B) 5 dps *pDR5::LUC* seedlings treated by NF and anchorene. Arrows in A and B indicate collets. (C) Effects of anchorene on 3 dps to 6 dps *pDR5rev::GFP* seedlings. White arrows indicate ANR initiation sites, and the red arrow indicates emerged ANR.

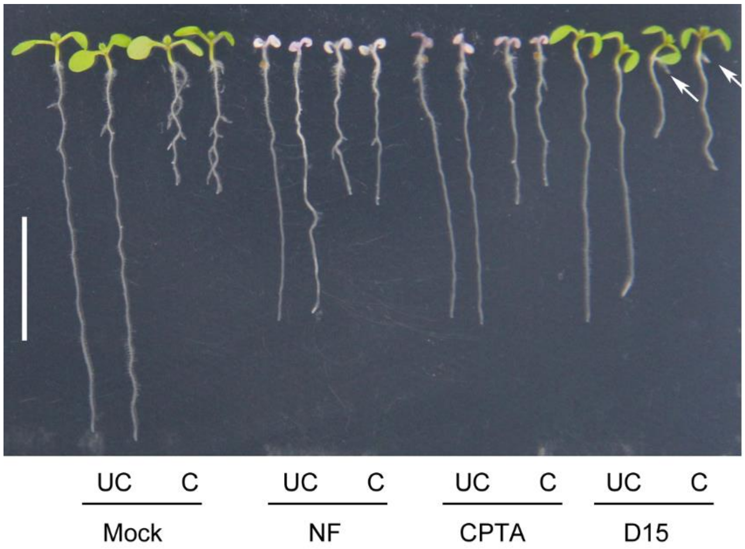


Fig. S5. Representative seedlings show the effect of different carotenoids biosynthesis inhibitors, NF and CPTA, and apocarotenoid inhibitor D15 on ANR and lateral root (LR) formation.

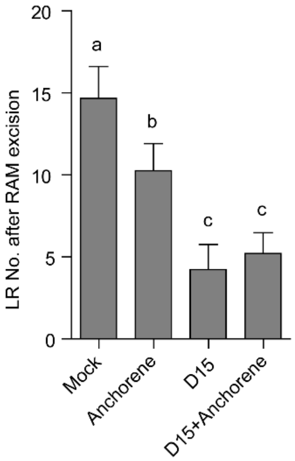


Fig. S6. The effect of anchorene and D15 on LR capacity (LR No. after RAM excision). Data were presented as mean ± SD (one-way ANOVA with Tukey multiple-comparison test, $P < 0.05$); different letters denote significant differences. 20 μM anchorene and 100 μM D15 were used.

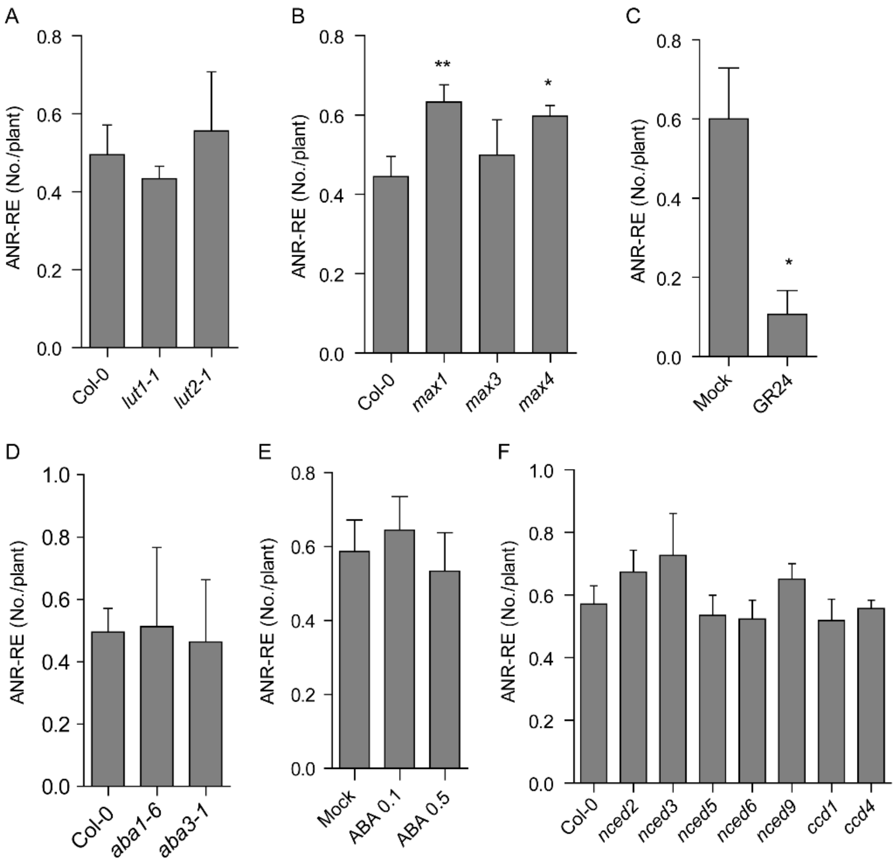


Fig. S7. Quantification of ANR formation after RAM excision (ANR-RE) from different carotenoids biosynthesis mutants (A), strigolactone biosynthesis deficient mutants (B), ABA biosynthesis deficient mutants (D) and carotenoids cleavage dioxygenase *ccd*/*nced* mutants (F). Effects of strigolactone analog GR24 (C) and ABA (E) on ANR-RE. Two-tailed Student's *t*-test was used for all statistical analysis, *$P < 0.05$, **$P < 0.01$. In A, B, D and F, data are presented as mean ± SD from 3 independent replicates. In C and E, data are presented as mean ± SE from 1 representative experiment; in C, n= 25, 28 respectively, 1 μM GR24 was used; in E, n=46, 45, 30 respectively, 0.1 and 0.5 μM ABA were used.

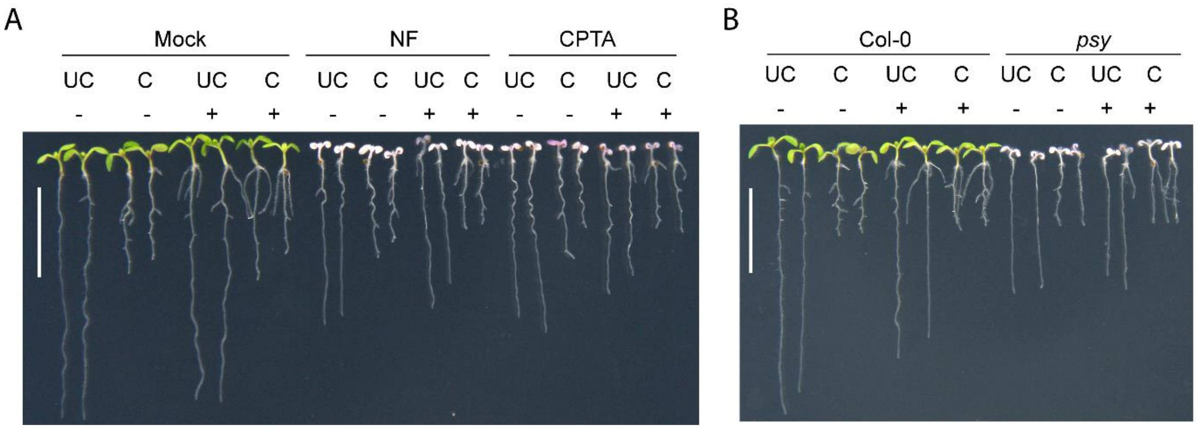


Fig. S8. Anchorene can restore the ANR-RE in carotenoid deficient seedlings. (A) Representative seedlings show the rescue of ANR-RE in NF- and CPTA-treated seedlings by anchorene. (B) Representative seedlings show that anchorene rescues ANR-RE in the *psy* mutant. In A and B, the symbols "-" and "+" indicate the absence or presence of anchorene application, respectively; "UC" indicates uncut RAMs, while "C" indicates cut RAMs; the scale bars are 1 cm.

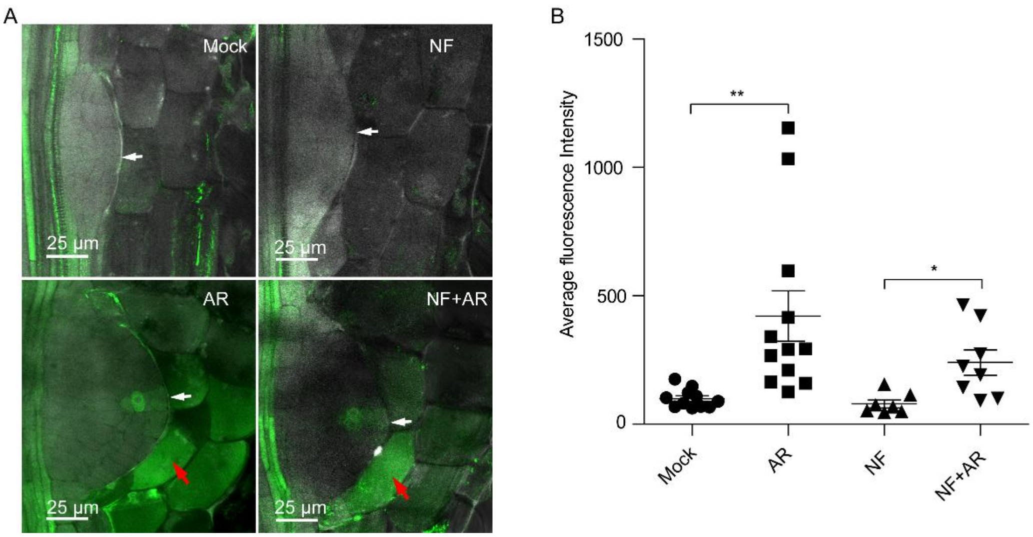


Fig. S9. Anchorene promotes DR5 expression in ANR primordia. (A) Anchorene activates the DR5 signal in ANR primordia and its abutting endodermis region of Mock and NF treated seedlings. White arrows indicate ANR primordia, and the red arrows indicate endodermis cells. *pDR5rev::GFP* was used as marker line. (B) GFP fluorescence intensity quantification in ANR primordia of different treated seedlings in A. data were presented as mean ± SD (two-tailed Student *t*-test, **P* <0.05 ***P* <0.01); n=11, 12, 7, 8 respectively from two independent experiments. 1 μM NF and 20 μM anchorene were used.

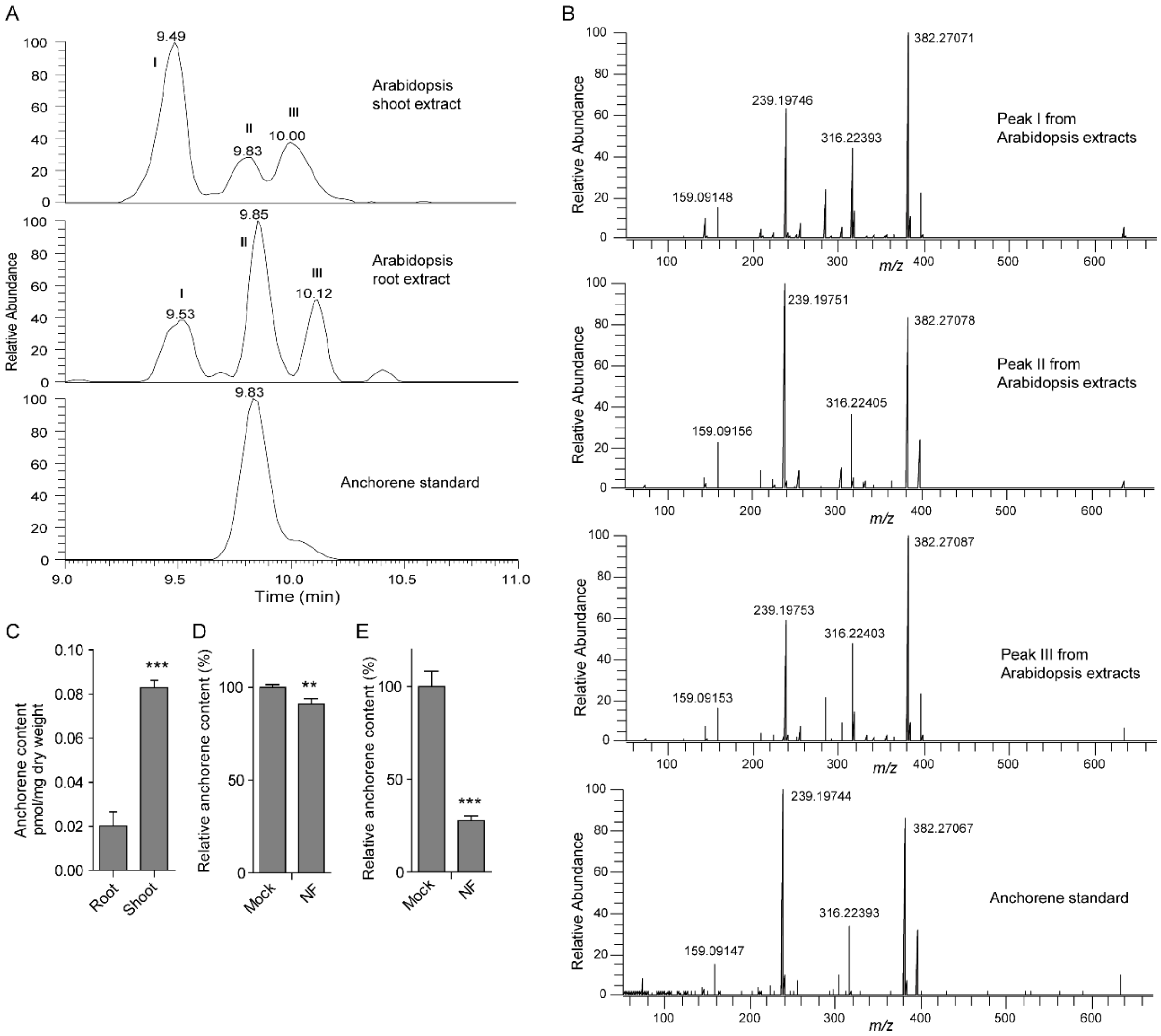


Fig. S10. The identification of potential anchorene isomers and the distribution of anchorene in Arabidopsis. (A) EICs of anchorene (peak II) and its isomers (peak I and peak III) from Arabidopsis shoot tissue (upper) and root tissue (middle), and EIC of authentic anchorene standard (bottom). (B) Product ion spectra of endogenous anchorene isomers including peak I, peak II and peak III from Arabidopsis extracts, and authentic anchorene standard. (C), Quantification of tissue-specified endogenous anchorene content in Arabidopsis. Relative anchorene content in mock and short term (D) or continuous (E) NF treated Arabidopsis seedlings. Two-tailed Student's *t*-test, in C and E, n=4; in D, n=3; $^{**}P < 0.01$; $^{***}P < 0.001$. 12-day-old seedlings were used for anchorene identification and quantification; in D, 11 day-old seedlings were treated by NF for another 24 hours. 2 μM NF was used.

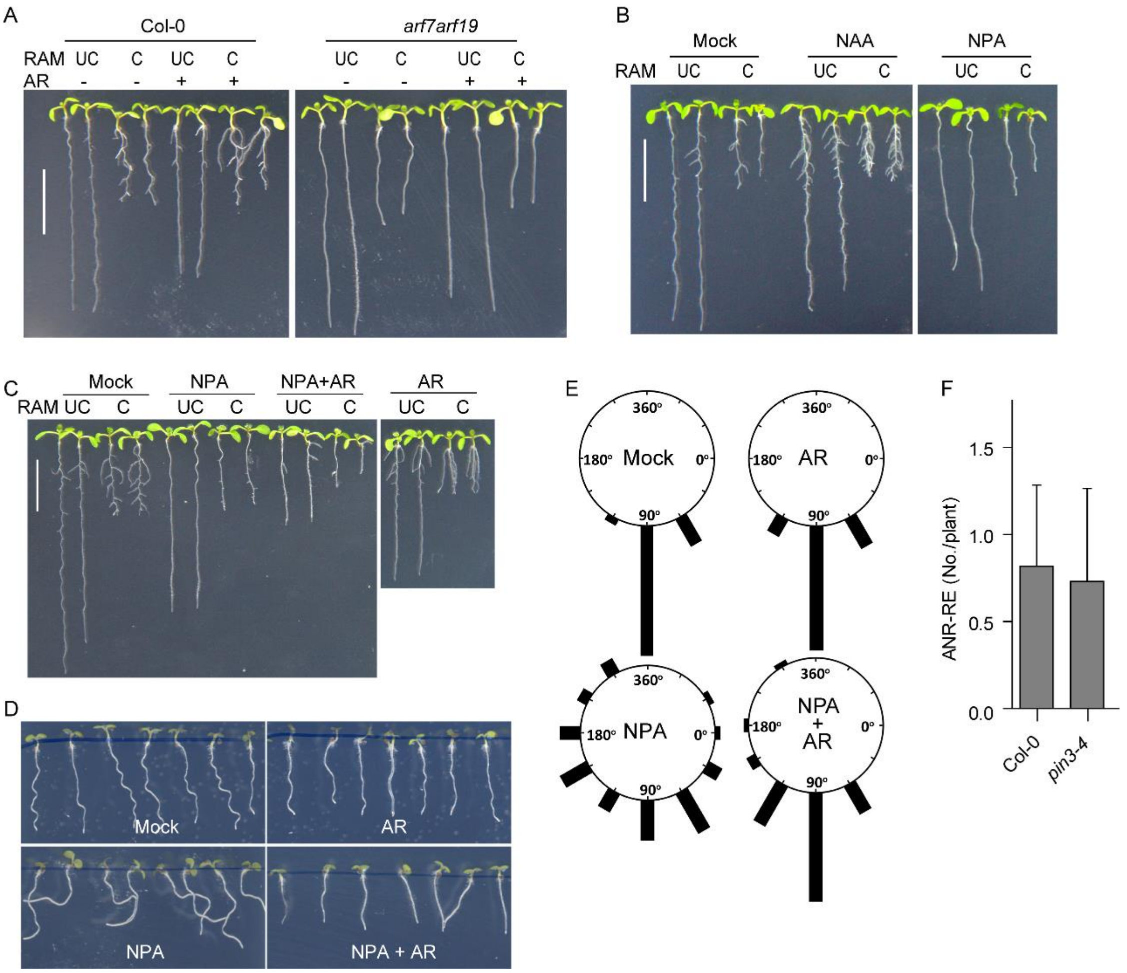


Fig. S11. Both auxin signaling and auxin distribution are important for ANR development. (A) Representative seedlings show the effects of anchorene on ANR formation in Col-0 and *arf7arf19*. (B) Representative seedlings show the effects of auxin analog NAA and auxin transport inhibitor NPA on ANR formation. (C) Representative seedlings show that anchorene partially rescue the inhibitory effects of NPA on ANR-RE. (D) Representative seedlings show that anchorene partially rescue the gravitropism loss of NPA treated seedlings. 5 dps seedlings were used for gravitropism analysis. ). (E) Anchorene partially rescued root gravitropism loss in NPA treated seedlings (n=27, 43, 41, 45 individually). The orientation of root growth was measured and then was assigned to one of twelve 30° sectors; the length of each bar represents the percentage of seedlings which displayed root growth within that sector. (F) ANR-RE of auxin transporter mutant *pin3-4* has no significant difference from corresponding wild-type seedlings. Two-tailed Student's *t*-test, n=26, 22 respectively. Scale bar, 1cm.

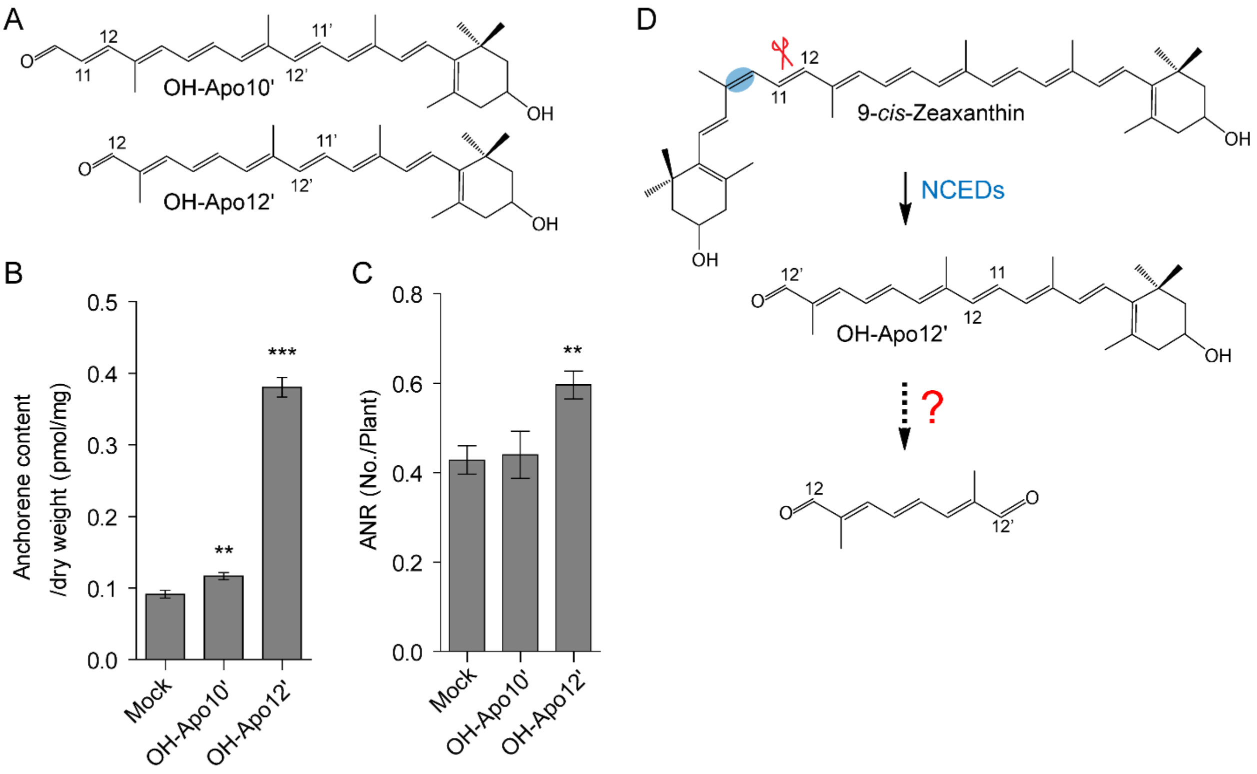


Fig. S12. OH-Apo12' is converted into anchorene in plants. (A) Structures of OH-Apo10' and OH-Apo12'. (B) Anchorene quantification in mock, OH-Apo10' and OH-Apo12' feeding Arabidopsis seedlings. (C) ANR emergence in OH-Apo10' and OH-Apo12' seedlings. Two-tailed Student's *t*-test, ***P* <0.01, ****P* <0.001; in B, n=4; in C data are presented as mean ± SD from 3 independent replicates; in B, 12-day-old seedlings were incubated with indicated chemicals for 6 hours and 20 μM OH-Apo10' and OH-Apo12' were used; in C, 10 μM OH-Apo10' and OH-Apo12' were used and the ANR No. was counted at 10 DAG. (D) Proposed formation of anchorene form 9-*cis*-Zeaxanthin. One of maize NCED, VP14, was shown to cleave 9-*cis*-Zeaxanthin (19).

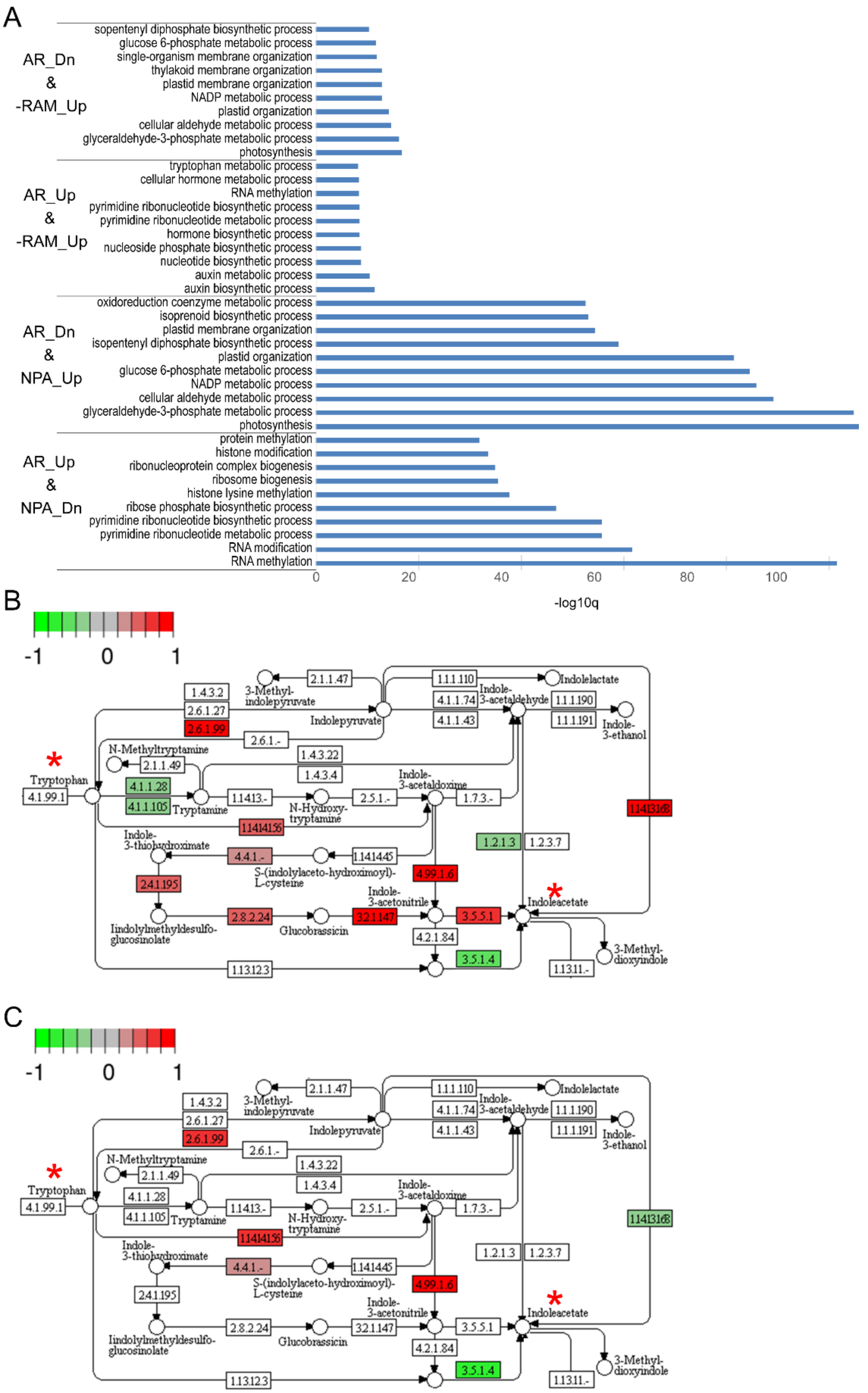


Fig. S13. RNA-Seq analysis of genes related to anchor root development. (A) Biological Processes (BP) analysis of overlapping genes of anchorene (AR) treatment with NPA treatment, and with RAM excision (-RAM). “Dn”

indicates down regulated genes, and “Up” indicates up regulated genes. Partial KEGG Arabidopsis tryptophan metabolism pathway (KEGG id: ath00380) to show the tryptophan-auxin biosynthesis pathway that regulated by anchorene (B) or RAM excision (C). The up-regulated genes and down-regulated genes are marked as red and blue colors, respectively; tryptophan and indole acetate were marked by red stars.

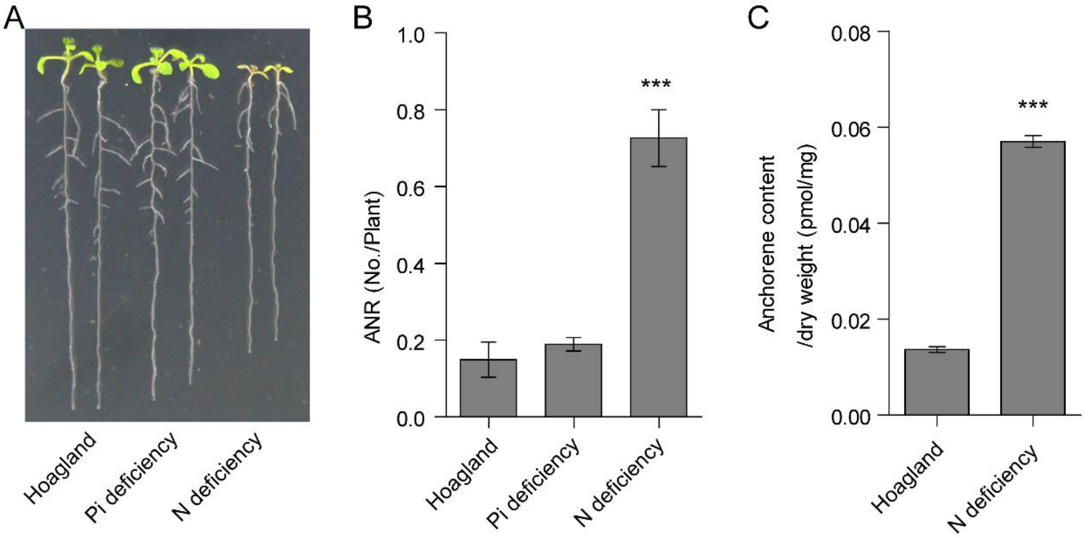


Fig. S14. Nitrogen (N) deficiency promotes anchor root formation. (A) Representative 10 dps Col-0 seedlings grown in agar plates with hoagland, phosphorus deficiency or N deficiency media plates. (B) Anchor root (ANR) No. of 10 dps Col-0 seedlings grown in agar plates with hoagland, phosphorus deficiency or N deficiency media plates. (C) Anchorene contents of the root tissues of seedlings grown in Hoagland and N deficiency media plates. 12 dps seedlings were uses. In B and C data are presented as mean ± SD; two-tailed Student’s *t*-test, ****P* <0.001; in B, n=4; in C, n=6, 3 respectively.

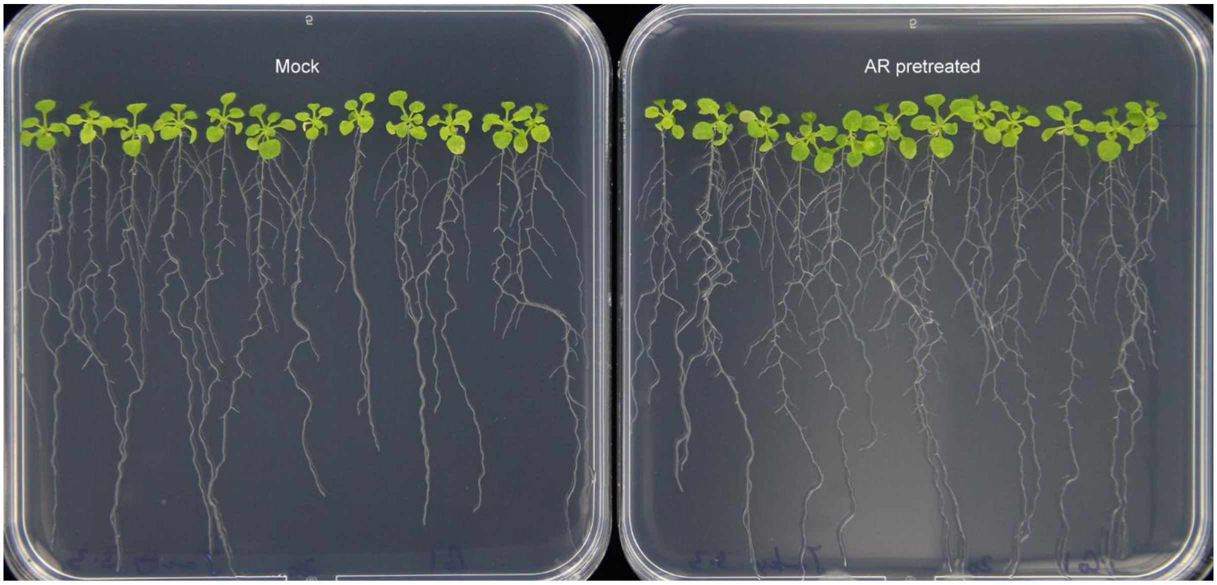


Fig. S15. Anchorene pretreatment increased the seedling biomass in Arabidopsis. 17-day-old seedlings grown in half MS agar plates with or without anchorene pretreatment are shown.

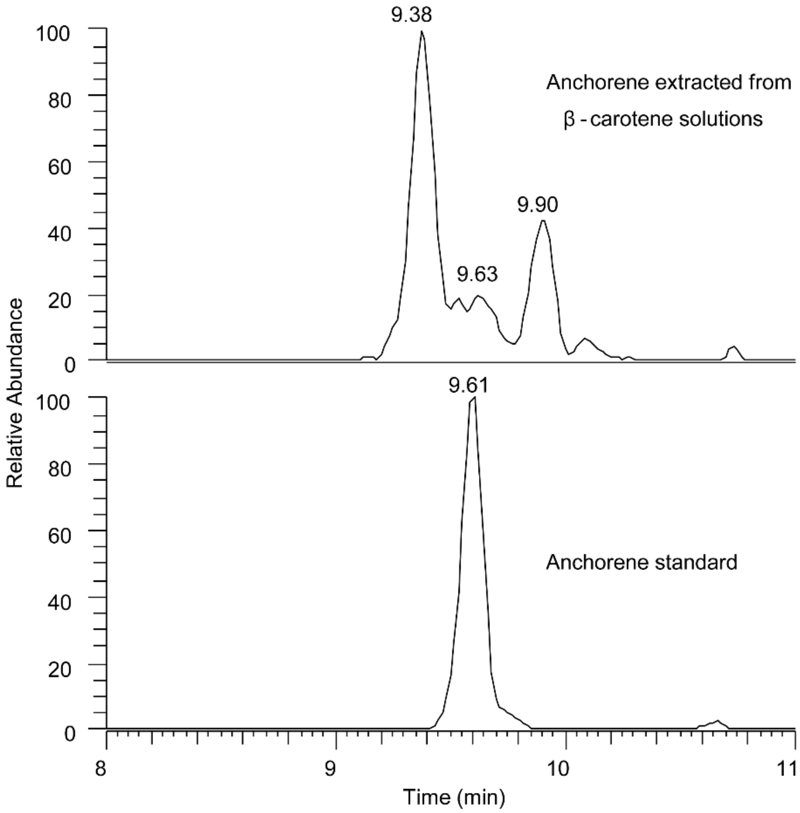


Fig. S16. Anchorene detection from a β-carotene organic solution. (A) EICs of anchorene extracted from β-carotene solution (upper) and and EIC of authentic anchorene standard (bottom). The solution was prepared by solving 10 µg β-carotene in 1 mL acetonitrile, followed by sonication for 20 min. Extraction, derivatization and LC-MS analysis were performed following the protocol used for plant material (s. below).

Table S1. The nutrient element composition in Argo soil and Silver sand.

| Element | mg/Kg | |
|---|---|---|
| | Argo soil | Silver sand |
| Fe | 780.53±2.19 | 1890.26±6.28 |
| K | 4001.53±82.19 | 758.83±24.47 |
| Mg | 11139.33±138.88 | 214.63±6.89 |
| Mn | 51.81±0.62 | 15.13±1.30 |
| P | 1468.06±25.08 | 25.57±2.98 |
| Zn | 58.20±1.75 | 17.77±0.42 |
| | g/Kg | |
| C | 439.93±8.22 | 3.56±0.34 |
| N | 10.69±0.54 | ND |
| S | ND | ND |
| H | 53.07±0.58 | 0.33±0.01 |

"ND" indicates non-detected.

Table S2. The pH value of Argo soil and Silver sand.

| | Argo soil | Silver sand |
|---|---|---|
| pH value | 6.37±0.03 | 7.05±0.03 |
